# A Preparation-Free Mixture-of-Experts Framework for Protein-Ligand Affinity Prediction

**DOI:** 10.64898/2026.07.24.740495

**Authors:** Huiming Bao, Shouliang Dong

## Abstract

Protein-ligand affinity (PLA) prediction is central to AI-driven drug discovery, but precise interaction-based methods require costly conformation preparation and data encoding, limiting their throughput. To reconcile accuracy with efficiency, we first investigate whether pre-trained molecular representation models can replace complex encoders. A unified and diverse assessment of sequence-, graph-, and image-based representations reveals both strong overall performance and family-wise variability, delivering the first practical guidance for encoder selection in PLA tasks. Next, to achieve high computational efficiency without sacrificing expressiveness, we adopt the mixture-of-experts (MoE) strategy from large language models. Systematic ablation studies uncover key design principles for deploying MoE in molecular prediction. The resulting model, HydrAffinity, is an interaction-free, dynamic sparse model that uses pre-trained encoders and MoE for parameter-efficient learning. It outperforms all interaction-free methods and matches state-of-the-art interaction-based methods on CASF-2016. Routing analysis confirms that MoE develops distinct, family-specific activation patterns, providing interpretable evidence of dynamic parameterization across protein classes. Zero-shot tests on DUDE-Z and LIT-PCBA further show strong EF^5%^ performance, making HydrAffinity a practical, scalable solution acting as an effective early-stage pre-filter.

**Graphic abstract:** 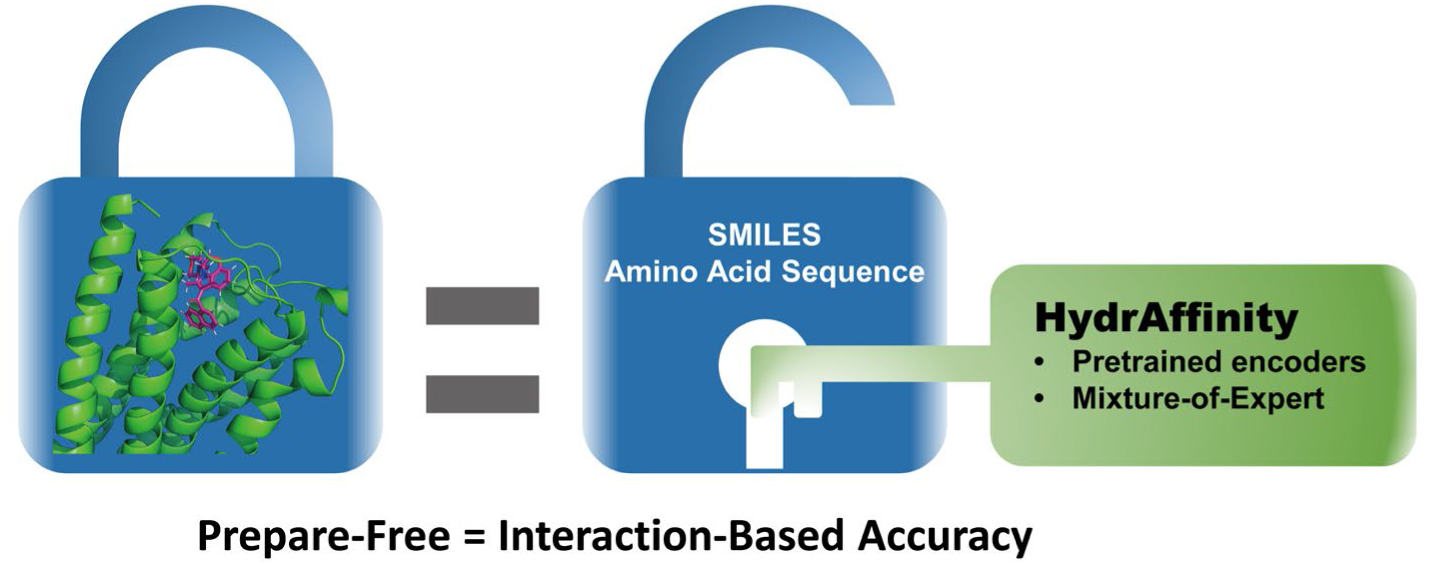

## Introduction

In the early stages of drug development, protein-ligand affinity (e.g., pK_d_) serves as a core metric for evaluating candidate compound activity. As chemical libraries expand exponentially, conventional wet-lab assays become prohibitively time-consuming and costly, rendering them impractical for large-scale screening^1^. In recent years, machine-learning–based affinity prediction models have emerged as a pivotal technology, enabling direct estimation of drug–target affinity^2^. These models generally fall into two paradigms: interaction-based and interaction-free^3^. Interaction-based approaches extract binding conformation and interaction information via molecular docking, graph neural networks (GNNs), or molecule interaction detection techniques, achieving higher accuracy by capturing atom-level interactions^4, 5^. By contrast, interaction-free methods simplify the input by relying solely on ligand chemical fingerprints (e.g., SMILES) and receptor amino-acid sequences, thereby omitting the docking step and offering superior scalability for large-scale, prepare-free virtual screening^6, 7^.

Although interaction-based methods retain an accuracy advantage, they suffer from high computational complexity and rely on high-quality docking results, which limits their practical deployment. Interaction-free methods, while computationally efficient, suffer from markedly lower prediction accuracy due to the absence of explicit atomic interaction modeling^8^. Three key challenges hinder the advancement of interaction-free models. First, a wide range of pre-trained molecular representation models exist (e.g., sequence-, graph-, and image-based), yet their performance on the PLA task remains largely unevaluated, leaving uncertainty about which representation and encoder is most suitable. Second, achieving competitive accuracy without explicit atomic interactions requires effective exploitation of implicit structural information embedded in molecular representations. Third, conventional Transformer layers that rely solely on self-attention or cross-attention mechanisms fail to replicate the fine-grained message passing inherent to GNNs, limiting the model’s ability to capture relational information between ligands and proteins.

To address the first challenge, we systematically evaluate diverse pre-trained models. Transformer-based ChemBERTa^9, 10^ and the ESM series^11, 12^ use a masked language model (MLM) training strategy, first converting molecules into linear textual representations before learning within a large-language-model (LLM) framework. In contrast, GNN-driven UniMol^13^ and ProSST^14^ focus on capturing molecular space features through graph structures. Each model offers distinct advantages in terms of input modalities—sequence^15, 16^, graph^17, 18^, or image^19, 20^—and architectural design, which effectively enhances the diversity of feature representations for PLA tasks.

To overcome the last two challenges, we explore architectural innovations. Conventional PLA approaches typically employ complex self-attention and interaction-attention mechanisms^21^, leading to a rapid increase in model parameters and exacerbating overfitting. Recently, a method based on a single Transformer module for multi-modal data processing has been proposed, significantly reducing architectural complexity^22^. More importantly, the mixture-of-experts (MoE) technique, when integrated into feed-forward networks (FFN), has been shown to achieve both lower computational cost and higher expressive power through sparse activation^23^. MoE has yielded remarkable performance gains in large language models and multi-modal architectures^24, 25^, opening up new possibilities for improving interaction-free affinity prediction.

In this study, we address all three challenges. First, we construct a comprehensive evaluation framework encompassing sequence-, graph-, and image-based molecular representations and systematically benchmark a suite of pre-trained models on PLA prediction. This analysis not only identifies the top-performing encoders but also reveals family-wise performance variations, providing the first practical guidance for encoder selection in protein–ligand affinity modeling. Second, we introduce a cascaded MoE architecture into the modality encoders, the multi-modal fusion transformer, and the affinity predictor, enabling effective exploitation of implicit structural information and surpassing the limitations of conventional attention architecture. The resulting model, HydrAffinity, is an interaction-free, dynamic sparse model that outperforms all existing interaction-free methods and achieves accuracy comparable to state-of-the-art interaction-based models on CASF-2016^26^—without any docking or 3D conformation preparation. Routing activation analysis reveals that MoE assigns distinct patterns to different protein families, confirming dynamic parameterization capability. Zero-shot evaluations on DUDE-Z^27^ and LIT-PCBA^28^ further demonstrate strong ranking performance compared with other methods. Collectively, this work demonstrates that interaction-free models can be both efficient and accurate, making them suitable for large-scale drug discovery.

## Methods

### Dataset

#### Data partition

This study adopts the standard partition established by EHIGN^3^, with data sourced from PDBbind v2016 (training set of 11,904 cases, validation set of 1,000 cases). The test set includes CASF-2013 (107 cases), CASF-2016 (285 cases), and the retained set from PDBbind 2019. It is important to note that there is no overlap among the training, validation, and test sets, ensuring a rigorous evaluation of model generalization. Due to varying requirements of pre-training models for molecular representations, there are minor discrepancies in data counts across datasets except for CASF-2016 (which includes CASF-2013) (detailed in Table S1).

#### Ligand representation

As shown in Table S2, we utilized three types of molecular representations for ligands: linear text descriptors (SMILES and SELFIELS), molecular structures (Graph), and molecular images (RDKit-generated 2D molecular images and PyMOL-generated^29^ ball, stick, and surface images). Detailed information about these models is described in Supplementary Section S3. For all these models, the maximum number of tokens or atoms was limited to 256. The image size was scaled to 224 or 512.

#### Receptor representation

We employed both sequence-only and structure-inclusive pre-trained models. Sequence-only models included ESM2^11^ (650M and 1.3B) and ESM3^12^ (esm3-open). Dual-modality models combining sequences and structures included SaProt^30^ (a pre-trained model using residue-level microstructures) and ProSST-2048^14^ (a pre-trained model using residue-level k-nearest neighbor subgraphs as structural information). We adopted a minimal-scale generation strategy for protein data: inputting each chain of the protein into the model separately, averaging the results across sequence, and further averaging across chains. This strategy aims to maximize the information of ligands and reduce computational burden in large-scale virtual screening.

### Mixture-of-Experts (MoE) architecture

#### Implementation of MoE modules

The detailed mathematical formula description of MoE is provided in Supplementary Section S4. In brief, we adopted the MoE implementation from large language models, e.g., DeepSeek V3^24^, including the router, shared experts, and routed experts. The router is a simple linear layer that projects inputs into expert activation scores.

To balance the activation among the expert networks, we incorporated noise routing, which functions only during training. Additionally, to improve computational efficiency, we implemented a concatenation-based MoE forward strategy. Unlike the common sample-wise loop, this strategy reduces the number of expert forward passes from *B* × *k* (where *B* is the batch size and *k* is the number of activated experts per sample) to the total number of experts *N*, which is typically much smaller than the batch size.

Regarding load balancing in MoE, our considerations focused on its impact on model expressivity and parameter utilization. Instead of employing auxiliary-loss-free balancing strategies (e.g., the bias-adjustment approach in DeepSeek-V3), we adopted a controllable cross-entropy-based constraint. Specifically, we apply a cross-entropy loss between the expert activation scores and a uniform target (all-ones vector), and multiply it by a scaling factor. Independent loss scales ranging from 1 × 10^−7^ to 0.05 were employed to flexibly control the intensity of this constraint. **Implementation of MoE modality encoders and MoE predictors.** Motivated by a recent study^31^ on routing strategy in large language models and the sequential nature of ligand modalities, we designed two routing strategies for the MoE modality encoder to investigate the effect of routing granularity: molecule-level routing, which treats each molecule as a single sample and routes it to selected experts, and token-level routing, which operates on individual tokens via tensor reshaping (detailed in Supplementary Section S4). The MoE predictor in HydrAffinity consists of multiple experts, each implemented as a standard MLP, forming an MoE module used as the affinity predictor.

### Implementation of mainstream MoE-FFN

For the mainstream MoE-FFN (Mixture-of-Experts Feed-Forward-Network), considering that the PLA task is relatively simple and the training data is limited, we set up only one shared expert with two options: the FFN layer used in ordinary transformers or the gated FFN used in large language model like DeepSeek V3. For routing experts, we configured 12 single-layer linear networks with an activation expert count of 1.

**Standard FFN**. Given an input vector **h** ∈ ℝ*^d^*^emb^, the standard FFN applies two linear transformations with an activation function *σ*(⋅) (SiLU in this paper):

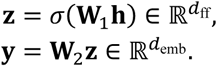

Where

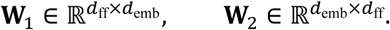

Dropout and activation functions are applied after *σ*(⋅), resulting in:

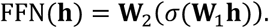

**Gated FFN**. Large language models like DeepSeek V3 incorporates a gating mechanism, multiplying hidden layer features with input features before mapping back to the input dimension. For **h** ∈ ℝ*^d^*^emb^, the computation is:

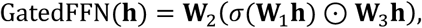

where

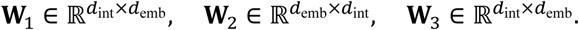

In this paper, W_1_ and W_2_ have dimensions (*d*_emb_, *d*_int_) and (*d*_int_, *d*_emb_), respectively, while W_3_ has dimension (*d*_emb_, *d*_int_).

The output of the gated FFN maintains the same dimensions as the standard FFN, ℝ*^d^*^emb^ 。

#### Computation

In the mainstream MoE-FFN, the routing expert ℛ consists of 12 single-layer linear networks:

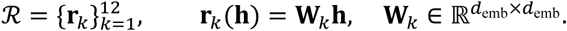

The shared expert ℰ uses a Gated FFN, with its output denoted as:

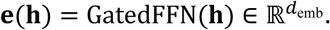

With an activation expert count of 1, each token is routed to a single expert. Let **g** ∈ ℝ^12^ be the normalized gating scores (unnormalized), and its softmax result is:

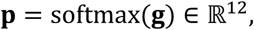

where 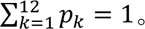

The MoE-FFN output is:

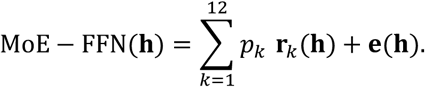

### Implementation of Hydraformer

For Hydraformer, it is designed as a modality-aware MoE-FFN, which expands the normalization layer and MoE-FFN layer to three times their original size compared to the mainstream MoE-FFN. It independently processes the CLS token (classification token, or called prediction token in this study), receptor modality, and ligand modality. More importantly, the three MoE-FFNs share a single shared expert. In this setup, the three modalities can freely interact through attention in the shared self-attention layer. Subsequently, in the three independent networks, the MoE-FFN leverages the advantages of dynamic sparse networks with high expressivity and low activation parameters while maintaining alignment across modalities via the shared large FFN.

Hydraformer extends the mainstream MoE-FFN by constructing independent MoE-FFNs for three modalities (CLS, receptor, ligand) while sharing a large Gated FFN as the shared expert. The computation process is as follows:

#### Modality Splitting

Given an input sequence **X** ∈ ℝ*^B^*^×*L*×*d*emb^ and modality index vector **m** ∈ {0,1,2}*^L^*, where:

*m_t_* = 0 indicates the CLS token,

*m_t_* = 1 indicates receptor modality,

*m_t_* = 2 indicates ligand modality.

For each modality *i* ∈ {0,1,2}, we define a mask:

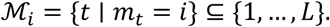

After the SDPA (shared multi-head self-attention) layer, the input H is processed by modality-specific normalization layers LN^(*i*)^_1_
, resulting in:

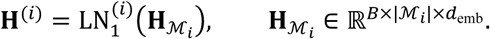

#### Independent MoE-FFN

Each modality *i* has 12 single-layer linear routing experts 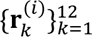, with an activation expert count of 2. The output of the routing experts is

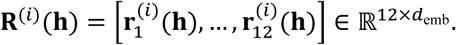

The gating mechanism G^(*i*)^ generates normalized gating scores **p**^(*i*)^ ∈ ℝ^12^ (softmax-normalized) and computes a weighted sum of the routing expert outputs to produce the activation expert output:

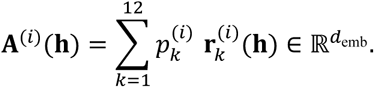

Subsequently, the shared expert ℰ (Gated FFN network) further transforms the original inputs for each modality to produce:

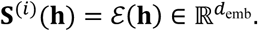

The final output of the MoE-FFN is given by:

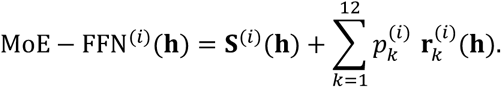

#### Shared Gated FFN Network

The three-way MoE-FFNs in Hydraformer share the same Gated FFN network as the shared expert. This shared expert is added after all modalities’ MoE-FFN outputs, forming a unified aligned representation. Specifically:

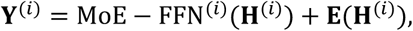

where E(⋅) = GatedFFNExpert(⋅) represents the output of the shared expert. Subsequently, each modality undergoes independent layer normalization LN^(*i*)^ and dropout processing to ensure residual connections and normalization remain independent across modalities.

### Training Objective and Load Balancing

The total load balancing loss for Hydraformer is the sum of the balancing losses from all three MoE-FFNs plus that of the shared expert. The overall training objective is:

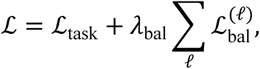

where *λ*_bal_ ∈ [10^−7^, 0.05] is the loss scale, and *l* indexes all MoE modules, ℒ_task_ is mean square error (MSE) loss of the prediction and the label.

### Model training

The model performance is evaluated by the root-mean-square error (RMSE), mean absolute error (MAE), Pearson’s correlation coefficient (R_p_), and the concordance index (CI). All experiments were repeated under three different fixed random seeds to ensure reproducibility and to obtain robust performance statistics (reported as mean ± std). All models were trained using the AdamW optimizer with a learning rate of 1 × 10^−4^ and weight decay set to 1 × 10^−6^. The batch size during training was either 196 or 256, and overfitting was controlled using early stopping: models were trained for a minimum of 300 epochs with a patience value of 100 epochs (in contrast, SOTA models employed a patience value of 500 epochs). Most experiments were conducted on an NVIDIA RTX 3080 GPU (10 GB VRAM); when memory requirements exceeded 10 GB, experiments were transferred to an NVIDIA RTX 5090 GPU (32 GB VRAM) for execution.

### Virtual Screening Benchmarks

Two public benchmarks were used to evaluate HydrAffinity’s screening power (ability to distinguish true binders from non-binders).

DUDE-Z^27^, the upgrade of DUD-E, provides three complementary subsets for 43 protein targets with 2,312 unique ligands: (i) DUDE-Z contains property-matched decoys (69,904, ratio ∼1:30); (ii) Extrema (732,309 decoys) introducing charge extremes (net charges −2 to +2) within ligand-like MW and cLogP; and (iii) Goldilocks (1,145,472 decoys) representing lead-like property space of ultra-large libraries. We evaluated HydrAffinity in four configurations: against the merged union of all three decoy subsets and against each subset separately. Because we only used the 1pt0LD ligand type from DUDE-Z, when a subset (Extrema or Goldilocks) lacked either actives or decoys, the missing ones were supplemented from the property-matched decoy set.

LIT-PCBA^28^ comprises 15 targets, 7,955 actives, and 2,644,022 inactives (overall ratio ∼1:332) from confirmatory dose-response PubChem BioAssays, with training/validation sets debiased by asymmetric validation embedding (AVE). For each target, we selected the highest-resolution crystal structure following PLANET’s strategy^32^, and performed the benchmark in both the training and validation set.

Due to extreme class imbalance and known data leakage in LIT-PCBA^33^, we adopted a strict zero-shot evaluation protocol on all experiments: the model was trained exclusively on the PDBbind set and directly applied to the target benchmarks without any fine-tuning, hyperparameter adaptation, or exposure to target-domain labels. Performance was measured by EF1%, EF5%, AUROC, and BEDROC (α = 80.5). EF at top *x*% of the ranked list is defined as:

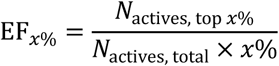

where: *N*_actives, top *x*%_ is the number of true actives in the top *x*% of the ranked list, *N*_total, top *x*%_ is the total compounds in the top *x*% (i.e., *x*% of the full library), EF measures how many times more enriched the actives are in the top fraction compared to random selection.

## Results

### Unimodal performance evaluation of pretrained molecular representation models

To systematically evaluate the effectiveness of various molecular representations and their pretrained models in protein-ligand affinity (PLA) task, we present a comprehensive summary in Table S3 of molecular representations based on serialized text, graph structures, and images, along with the corresponding pretrained models. Following the dataset partitioning criteria established by EHIGN^3^, we generate vectorized representations of ligands and receptors from the PDBbind v2019 dataset^34, 35^ using each pretrained model. The example of each ligand representations is provided in Table S2. Subsequently, we visualize the resulting embeddings with UMAP; the results reveal that all pretrained models cluster molecules of the same class in nearby regions, forming clear cluster structures (Figure S1). This observation indicates that these models have successfully captured chemical or biological information about molecules in latent space.

Subsequently, we constructed a concise projector–Transformer–predictor framework (Figure 1a) and used this architecture to systematically evaluate the PLA performance of all pretrained models (Figure 1b, c. More performance results of scratch embedding are shown in Figure S2). Single-modal training first verifies the discriminative power of pretrained embeddings and their relevance to the PLA task, enabling rapid identification of high-potential modalities. Experimental results of CASF-2016 benchmark dataset^26^ show that ligand representations based on visual features lag significantly behind linear text and graph-based molecular representations in the PLA task, suggesting that visual features are insufficiently effective for molecular encoding.

**Figure 1.**
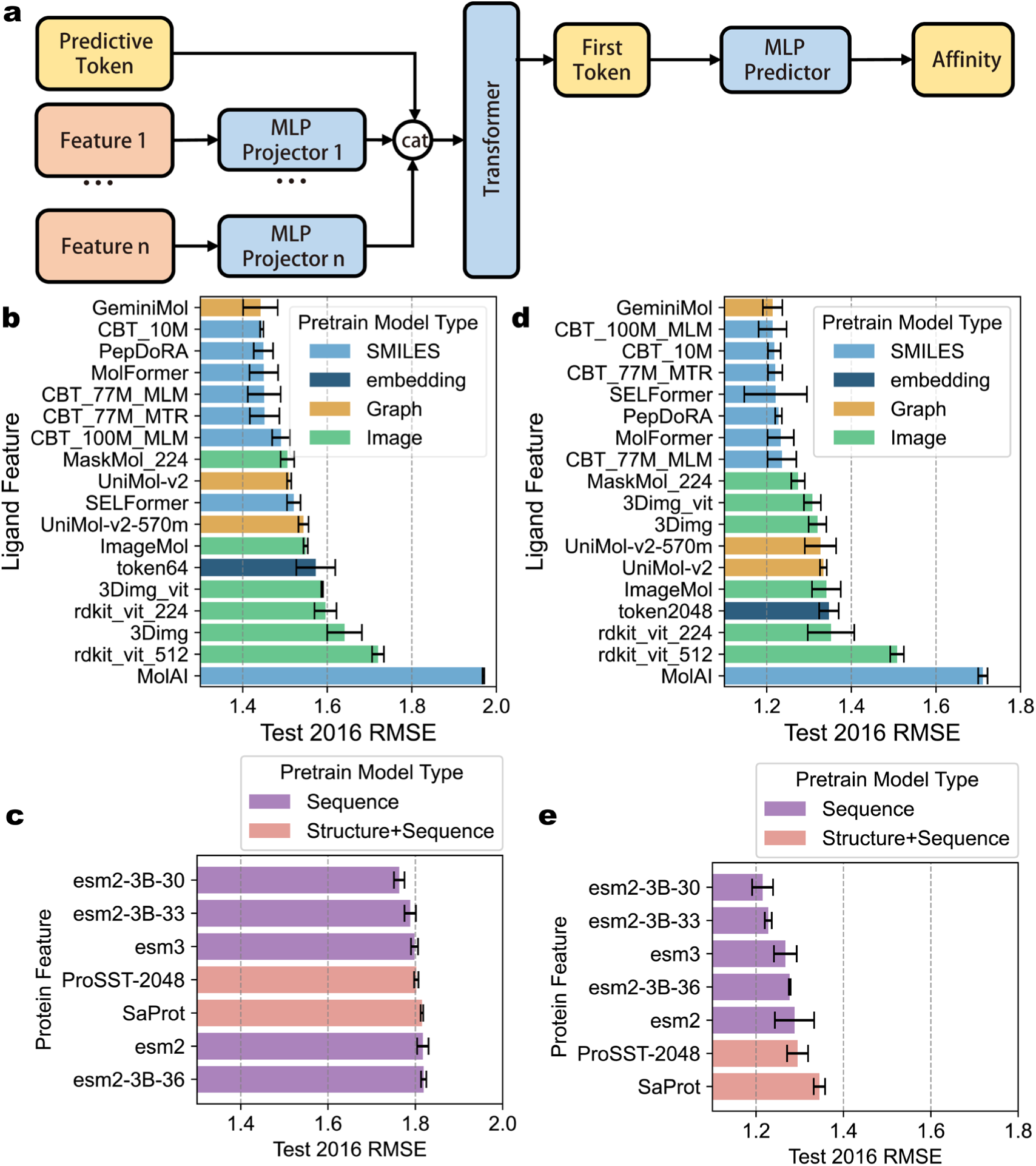
**a,** A Protein-Ligand-Affinity (PLA) task model architecture based on a multimodal Transformer. **b-c,** Unimodal results: (b) performance of the small-molecule ligand’s molecular pre-training feature embeddings, and (c) performance of the receptor protein’s pre-training feature embeddings, both evaluated on the PLA task. CBT refers to the abbreviation of ChemBERTa. **d-e,** Bimodal results: (d) performance of the small-molecule ligand’s molecular pre-training feature embeddings, and (e) performance of the receptor protein’s pre-training feature embeddings, both evaluated on the PLA task. Error bars on the bar charts represent standard deviation (std), with n = 3.

### Bimodal performance evaluation of pretrained molecular representation models

To further investigate the complementarity and conflicts among representations, we analyzed the intrinsic and extrinsic associations of the features. Figure S3 shows that there is a lack of complementarity among all receptor representations in PLA task as dual modal. Aligning the TM-score^36^ matrix within protein families with the similarity matrix of pre-trained features revealed that protein pre-training embeddings and family similarity exhibit no obvious correlation with performance on the PLA task (Figure S4), suggesting that the molecular semantics learned by pre-training models may not translate into superior PLA performance.

Based on the aforementioned findings, we selected the PepDoRA^37^ model—characterized by a sequence-representation dimension while maintaining a low embedding dimension—as the ligand benchmark. In parallel, owing to its superior performance, we adopted the esm2-3B-33^11^ model, which effectively balances coarse- and fine-grained feature representations, as the receptor benchmark.

Results of bimodal performance (Figure 1d, e) show that embeddings from SMILES-based pre-trained models consistently outperform graph- or image-based alternatives in PLA prediction. Graph-based pre-training exhibits higher performance variability, while image-based approaches yield the weakest results. Among image encoders, MaskMol^19^—an image model that models atomic and chemical bond information—achieves the best performance, highlighting that embedding chemically meaningful cues from pre-training is more critical than providing more details. For protein encoders, larger pre-training scales generally improve PLA prediction, yet the largest model (ESM-2 3B^11^) performs best at an intermediate layer rather than the final output. This further suggests that fine-grained representations carry greater value than coarse-grained ones, challenging the conventional preference for coarse abstraction in high-level tasks.

Notably, the dual-modal approach consistently outperforms single-modal methods, highlighting the importance of incorporating complete information for accurate PLA predictions. Furthermore, the shift in rankings—such as ChemBERTa_100M_MLM moving from a second-tier position in single-modal experiments to first-tier in dual-modal setups—underscores the complexity of modality interactions and the necessity of systematic evaluations to fully understand their impact on model performance.

### The MoE architecture design

In recent years, MoE has become increasingly prominent in large language models for its ability to scale model capacity efficiently^38, 39^. Given the multimodal nature and parameter sensitivity of preparation-free PLA prediction, we hypothesize that MoE could enhance model expressiveness while maintaining computational feasibility. Building on this, we incorporate MoE into the PLA algorithm.

First, we implemented MoE on the predictor. In the previous architecture (Figure 2a), the predictor was a simple two-layer multilayer perceptron (MLP). As shown in Figure 2b, we treated the original predictor as an expert module and stacked N experts under a single router. To balance expert utilization, we introduced a balance loss on the router’s activation scores, using cross-entropy against a uniform distribution. We also added noise before the router to encourage diverse expert activation and avoid parameter waste.

**Figure 2.**
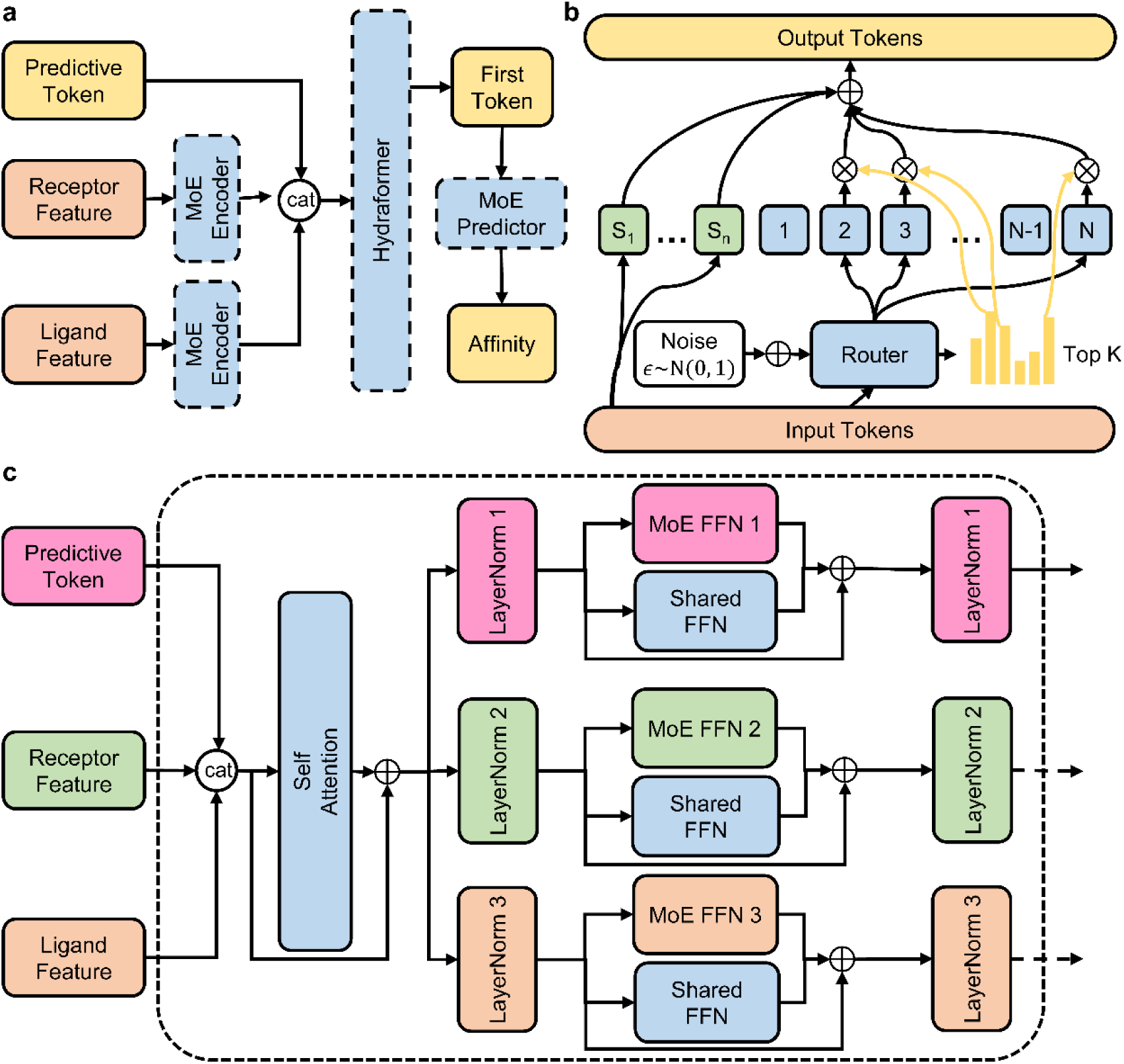
**a,** Schematic diagram of the HydrAffinity architecture. Modules within the dashed boxes are all MoE modules. **b,** Schematic diagram of the MoE module architecture. The module in the green box is the shared expert, and the modules numbered 1∼N in blue boxes are routing experts. The illustrated routing aggregation strategy is weighted. **c,** Schematic diagram of the Hydraformer architecture. Note that the final dashed arrow indicates that, only the predictive token contributes to the output in the last layer.

Experiments show that, in predictor MoE module, as the total number of experts increases, model performance first improves then deteriorates (Figure S5a), suggesting that smaller tasks require fewer experts. When more experts are activated, performance becomes unstable; this can be mitigated by router noise or a weighted-sum output strategy (Figure S5b). Moreover, router noise can enhance MoE performance, while the weighted-sum aggregation strategy effectively stabilizes MoE performance even under high expert activation ratios.

Next, we applied MoE to the modality encoder. Like the predictor, the modality encoder (currently a single-layer MLP) is treated as an expert module. Because the ligand modality contains sequential information, we incorporated a token-level tokenization scheme^31^ into the MoE; experiments demonstrate that token-level MoE markedly outperforms sequence-level MoE (Figure S6). This suggests that fine-grained, token-wise routing is better suited to capture the local structural variations in ligand sequences. In contrast to the ligand modality, the receptor MoE did not exhibit a clear performance trend with changes in the total number of experts or the activation ratio (Figure S7).

Finally, we introduced MoE into the modality-fusion layer—the Transformer—drawing inspiration from the Higgs Audio framework. We extended the mainstream MoE-FFN architecture into a multimodal version suitable for PLA tasks, termed Hydraformer, which utilizes a separate small-router expert group for each modality and shares a large, shared FFN across all MoE-FFNs for parameter sharing (Figure 2c). To reduce computational cost and achieve implicit modality alignment, we innovatively designed a “large-shared-experts + small-router” architecture: shared experts learn cross-modal common information, enabling implicit alignment; the small router captures modality-specific information while preserving computational savings. Additionally, we made LayerNorm modality-independent and processed multimodal inputs in parallel before separating them for modality-specific networks. Experimental results show that Hydraformer outperforms the conventional MoE-FFNs implementation (Figure S8), demonstrating that the combination of modality-specific small MoEs and large shared experts is more effective than a single multimodal MoE, as it better balances modality-specific adaptation with cross-modal common feature learning.

### Performance on PDBbind dataset

Combining the MoE modality encoder, MoE predictor, and Hydraformer, our model achieves an average RMSE of 1.161 on the CASF-2016 dataset, surpassing all interaction-based methods and reaching state-of-the-art performance of interaction-based method (Table 1). Additionally, the model attains a CI of 0.826 (0.006) on the same benchmark, indicating strong ranking capability (Table S4). All compared results are collected from public reports, and the core training-to-testing split (PDBbind to CASF-2016) remains consistent across studies: removing CASF-2016 items from training set to make no overlap between from training set and test sets. The minor protocol variations mainly involve different PDBbind versions (v2016 vs. v2020) and unavoidable instance losses during data encoding; a full comparison is provided in Supplementary Section S15. Moreover, these ablation results (Table 2) show how MoE can be tailored to multimodal architectures: weighted-sum aggregation stabilizes outputs, and a shared-expert design aligns modalities efficiently.

**Table 1.** Comparison results of the proposed HydrAffinity and baselines on CASF-2016 set.

| Method | Name | CASF-2016<br>RMSE | CASF-2016<br>R<sub>p</sub> |
| --- | --- | --- | --- |
| Interaction-Based | GEMS<sup>40</sup> | 1.223 | 0.832 |
|  | Surface-based<sup>41</sup> | 1.151 | 0.851 |
|  | EHIGN<sup>3</sup> | 1.150 (0.022) | 0.854 (0.004) |
|  | AGL-EAT-Score<sup>42</sup> | 1.150 | 0.873 |
|  | FGNN<sup>43</sup> | 1.140 | 0.873 |
|  | **Dynaformer<sup>44</sup>** | **1.114** | - |
| Interaction-Free | Multi-PLI<sup>45</sup> | 1.437 | 0.750 |
|  | PLMAM-PLA<sup>46</sup> | 1.223 | 0.838 |
|  | **HydrAffinity-S1 (ours)** | **1.162 (0.036)** | **0.846 (0.010)** |
|  | HydrAffinity-M3 (ours) | 1.170 (0.035) | 0.844 (0.011) |

Note that since CASF-2016 is widely used in the protein-ligand affinity studies and provides good comparative significance, this study primarily compares performance on the CASF-2016 dataset. The performance of all other algorithms is sourced from their reported data, presented as mean (std), n=3. HydrAffinity-S1 and HydrAffinity-M3 stand for bimodal architecture and multimodal architecture. Bold text indicates the best performance. A detailed comparison of dataset partition is provided in Table S6.

**Table 2.** Ablation study results of Hydraformer module on the 2013 core set, 2016 core set, and the 2019 holdout set.

| Name | 2013 RMSE | 2013 R<sub>p</sub> | 2016 RMSE | 2016 R<sub>p</sub> | 2019 RMSE | 2019 R<sub>p</sub> |
| --- | --- | --- | --- | --- | --- | --- |
| **HydrAffinity-S1 (Hydraformer)** | 1.356 (0.025) | 0.813 (0.009) | 1.162 (0.036) | 0.846 (0.010) | 1.457 (0.015) | 0.639 (0.003) |
| w/o weighted gatter (sum) | 1.317 (0.047) | 0.827 (0.014) | 1.172 (0.020) | 0.844 (0.003) | 1.474 (0.023) | 0.635 (0.003) |
| w/o Hydraformer (MoE-FFN) | 1.313 (0.046) | 0.829 (0.013) | 1.179 (0.025) | 0.842 (0.006) | 1.497 (0.026) | 0.628 (0.008) |
| w/o Hydraformer (Transformer) | 1.377 (0.032) | 0.809 (0.010) | 1.184 (0.020) | 0.839 (0.007) | 1.444 (0.048) | 0.640 (0.012) |
| w/o shared expert | 1.361 (0.052) | 0.812 (0.017) | 1.206 (0.050) | 0.832 (0.016) | 1.448 (0.008) | 0.630 (0.005) |

Note that 2013, 2016, 2019 stands for the CASF-2013, CASF-2016, PDBbind 2019 holdout set in this study. data is presented as mean (std), n=3.

To examine the generalizability of the findings regarding pretrained model evaluation and MoE design, we additionally evaluated our model on the Davis^47^, KIBA^48^, and BindingDB^49^ datasets, which are widely used by many interaction-free models^50–52^. Because these three datasets contain relatively few proteins and ligands, we directly adopted the best-performing ligand and receptor pretrained models identified in our earlier evaluation, and simplified the MoE component by retaining only the MoE predictor, in order to mitigate potential overfitting. Using the performance data collected by DeepDTAGen^50^, we found that our model achieves the best MSE among all compared methods, and the best CI on the Davis and KIBA datasets, while it does not show an advantage in *r*^2^ and AUPR (Supplementary Section 16). This indicates that the pretrained model selection results and the design principles of MoE identified in this study can be readily transferred to other datasets with satisfactory performance.

A critical yet challenging issue for interaction-free methods is the out-of-distribution (OOD) problem. To assess this, we retrained and evaluated HydrAffinity using the stricter PDBbind v2020 split proposed by GEMS^40^. According to their protocol, complexes with similar protein structures, ligands, or affinity labels to the test set are removed from the training set. Consequently, all reported models exhibit performance degradation under this split. We acknowledge that HydrAffinity also shows a clear drop in performance; nevertheless, it remains close to the top tier of interaction-based methods (Table S5). More importantly, the multimodal HydrAffinity outperforms its unimodal counterpart under the stricter split, yet it does not exhibit an advantage under general relevance-filtering conditions. This suggests that while multimodal information provides additional cues to overcome OOD challenge, it may sometimes also introduce data noise.

### Analysis of MoE modality-adaptive router

MoE is a highly dynamic architecture. To investigate its dynamic activation characteristics, we extracted the router activation scores from the aforementioned MoE experiments. We first examined activation bias, where the router tends to assign high scores to a few experts, causing their frequent activation (“expert overwork”). In extreme cases, less-activated experts receive scant data, becoming under-trained, which can degrade performance and lead to inefficient parameter utilization^24^.

By computing the coefficient of variation (CV) for MoE router activation score across multiple experimental runs, we found that for predictors, CV strongly correlates with the total number of experts (Figure 3a, *R*^2^ = 0.85) but not with the activated number of experts (Figure 3b, R^2^ = 0.02). The routing-aggregation method also affects the distribution: arithmetic summation yields more uniform activation, whereas weighted summation leads to dominance by few experts (Figure 3c). Critically, since CV does not correlate with model performance (Figure S9a, Figure S11b), we posit that for the PLA task, an appropriate total number of experts is sufficient. When this number increases, the model maintains efficiency by dynamically silencing a subset of experts, effectively compressing the usable parameter scale without harming performance (Figure S10).

**Figure 3.**
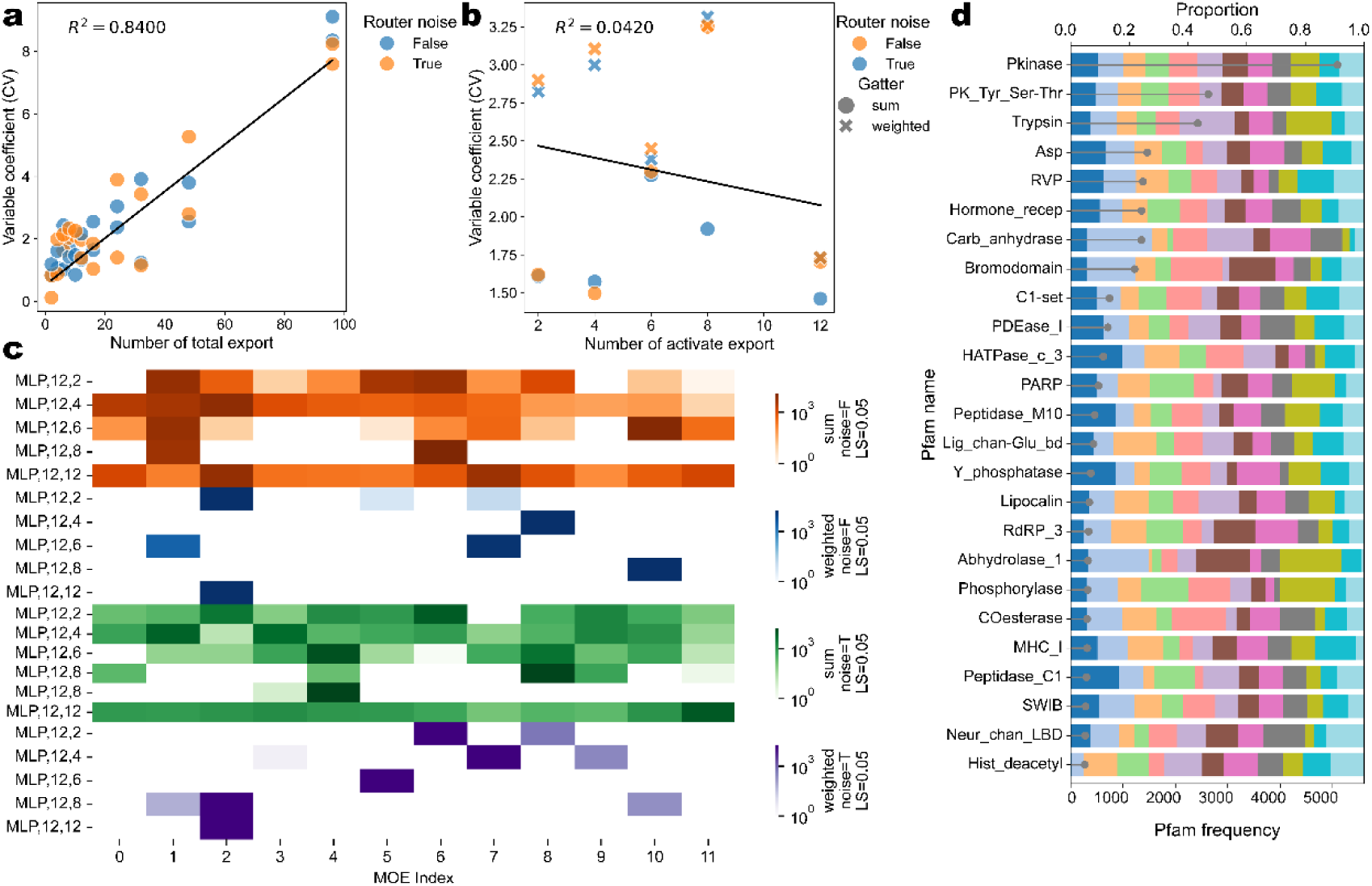
Relationship between the total number of experts (**a**), activated experts (**b**) and the coefficient of variation (CV) of the routing activation scores in the Predictor MoE. This panel includes experimental data for MoE balance loss scales of 0.05 and 1. **c,** Heatmap of expert activation patterns in the Predictor MoE. This panel counts the activation frequency of expert modules across all datasets. In the legend, weighted and sum represent routing aggregation strategies; noise represents the noise router; LS represents the MoE balance loss scale. **d,** Activation patterns of the Receptor MoE module for protein data from different families. This experiment uses a Receptor modality encoder with an MoE architecture only (total experts: 12, activated experts: 4, using router noise, using the sum routing aggregation strategy, CASF-2016 performance: 1.200 (0.032), mean (std)). The stacked bar chart shows the percentage of activation counts for all data in the dataset by the 12 experts (top x-axis). The lollipop chart shows the count of protein families in the dataset (bottom x-axis).

Building on this, we further analyzed the activation distribution of receptor-modality MoE routing across different protein families. To understand how routing relates to biological structure, we leveraged the protein family classifications (through HMMER^53^ and Pfam^54^) that underlie the semantically meaningful clusters in the pre-trained embeddings. Subsequently, we tabulated the activation of each expert by family. In larger families, activation bias was mild; as family size decreased, more families showed dominance by one or a few experts. Activation patterns differed substantially across families (Figure 3d), endowing the model with a degree of interpretable, target-family-aware specialization.

### Virtual screening capability evaluation

In real-world large-scale virtual screening, a single receptor is often confronted with millions to billions of candidate molecules, which may be generated by AI models or drawn from large chemical libraries. For an interaction-free PLA model, achieving acceptable enrichment with minimal input preparation is a challenging yet crucial goal. HydrAffinity is designed to address this need. However, strong performance on CASF-2016 does not guarantee effective virtual screening in practice. Therefore, we further evaluate HydrAffinity on two challenging benchmarks: DUDE-Z^27^ and LIT-PCBA^28^, under zero-shot settings (i.e., no retraining on any datasets).

DUDE-Z benchmark includes three subsets: DUDE-Z, Extrema, and Goldilocks, which represents different molecular groups. For the EF5% metric (enrichment factor at 5% of the ranked list), most models achieve AUROC around 0.7 and EF5% values between 20 and 40 on the DUDE-Z and Goldilocks subsets, indicating reasonable enrichment capability. However, all models perform worst on Extrema, with EF5% on Goldilocks up to four times that on Extrema for several models (Figure 4a). AUROC values on Extrema hover around 0.5 for all models, confirming that ligands with extreme charges severely disrupt all molecular modalities (Figure 4b). Sequence-based (SMILES, SELFIES), graph-based, and 2D image representations all encode charge information, and the distribution shift caused by extreme charges in Extrema leads to significant performance degradation. This issue can likely be mitigated by including similar charged molecules in the training set.

**Figure 4.**
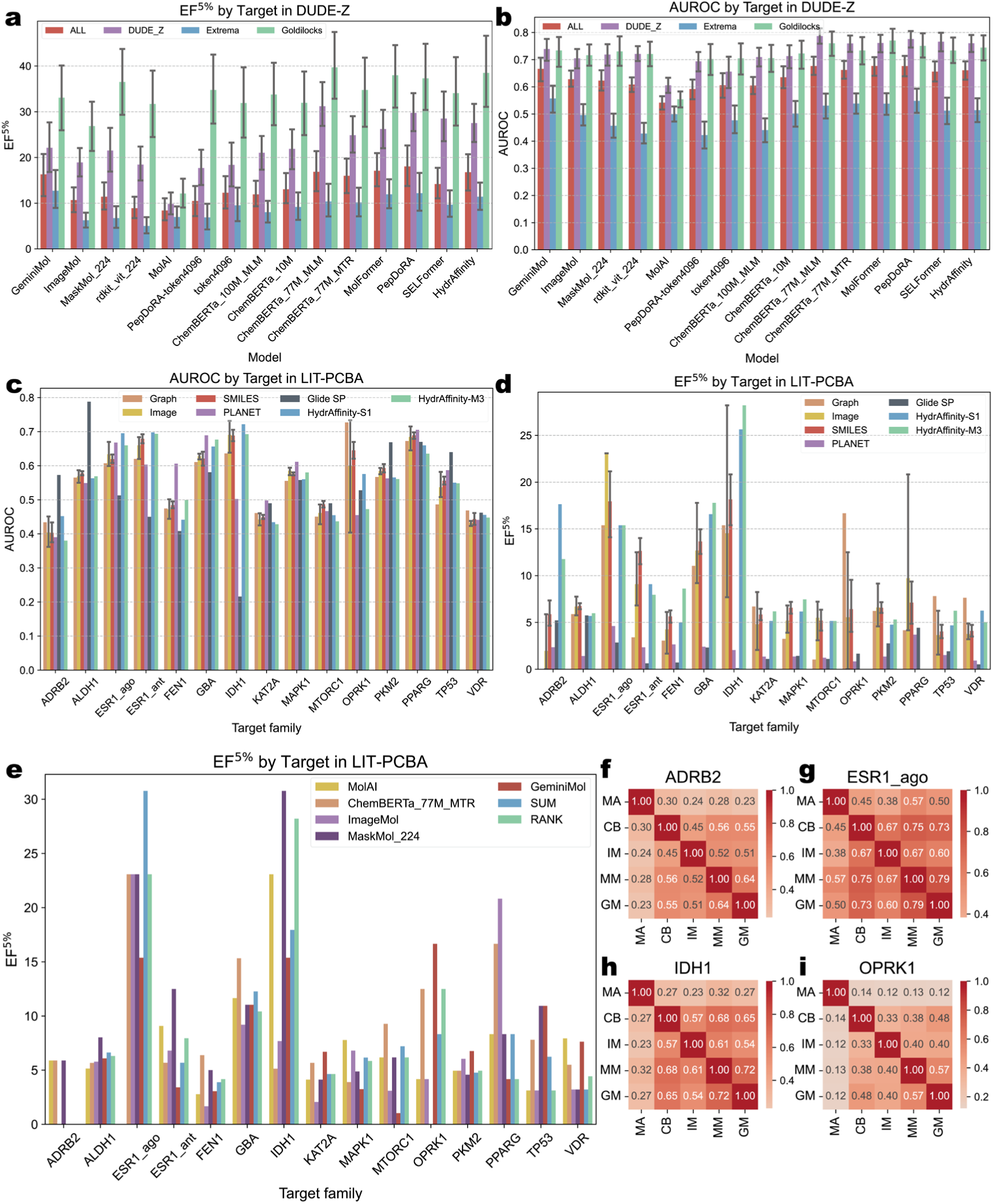
**a** (EF5%), **b** (AUROC) result of each molecular presentation pre-train model on the DUDE-Z dataset. **c** (AUROC), **d** (EF5%) result of each molecular presentation pre-train model on the LIT-PCBA dataset. The results of PLANET and Goldilocks are taken from the works of PLANET. HydrAffinity-S1 and HydrAffinity-M3 stands for bimodal architecture and multimodal architecture in Table 1. **e**, EF5% result of MolAI, ChemBERTa_77M_MTR, ImageMol, MaskMol, GeminiMol, summation of every model and average rank of every model. **f-I**, rank correlation of each model on four targets in LIT-PCBA. MA, CB, IM, MM and GM represent the abbreviations of the above five models in sequence. Error bars on the bar charts represent 95% confidence level.

LIT-PCBA poses a harder virtual screening challenge than DUDE-Z, featuring more decoys, highly imbalanced ratios, and fewer targets. To benchmark HydrAffinity, we compare it with PLANET^32^ (a GNN-based interaction-free method trained on PDBbind v2020) and the classical docking method Glide SP, all under zero-shot conditions. For EF1% and AUROC (Figure S11, Figure 4c), no clear differences are observed among the tested models; none consistently outperforms the others across multiple targets. However, for EF5%, HydrAffinity establishes a substantial advantage on most targets (Figure 4d). Since EF5% measures enrichment within a broader top fraction, this highlights HydrAffinity’s effectiveness as a preparation-free, rapid pre-filter for early-stage screening, rather than for final top-hit prioritization (where EF1% is more relevant). By retaining a larger fraction of active molecules (higher EF5%) while drastically shrinking the compound library, HydrAffinity enables subsequent, more accurate but conformation-demanding interaction-based methods to be applied efficiently on the reduced set for fine-grained enrichment.

Notably, different molecular representations and pre-trained models exhibit varying virtual screening performance. For example, MolAI (an LSTM-based model using SMILES input) shows lower average performance on all DUDE-Z subsets but achieves leading EF1% on the IDH1 and PPARG targets in LIT-PCBA (Figure S11, Figure 4d). GeminiMol (a graph-based model) delivers a large EF1% lead on the OPRK1 target but performs modestly on others (Figure 4e). These observations suggest that different pre-trained models have distinct molecular projection biases, making some more suitable for PLA tasks on certain molecular groups. As shown in the correlation matrix of affinity predictions on LIT-PCBA (Figure 4f-i), the rankings across models exhibit moderate consensus, yet no single model’s average prediction is consistently superior (Figure 4e). This indicates that multi-modal prediction has the potential to improve generalization, but simply averaging predictions from multiple models is not an effective strategy.

## Discussion

Many protein–ligand affinity algorithms employ pre-trained molecular representation models to extract information from molecules, a process that is substantially more efficient than interaction-based approaches. Through large-scale, systematic evaluation of diverse molecular representations and their pre-trained models, we identified several under-explored yet top-performing models. More importantly, our systematic evaluation reveals that Transformer-based SMILES embeddings consistently outperform graph- or image-based alternatives for PLA prediction, suggesting that large-scale language modeling of linear sequences captures chemically relevant patterns more effectively than spatial or visual encodings. The observation that the best protein representation comes from an intermediate layer of ESM-2 (rather than its final output) implies that fine-grained, hierarchical features are more informative than coarse-grained abstractions—a finding that may generalize to other biomolecular tasks.

The PLA algorithm suitable for virtual screening also needs to have excellent predictive ability and extremely high computing performance. Starting from architectural innovation, we developed HydrAffinity. Its cascaded MoE design addresses two key limitations of interaction-free methods: it leverages implicit structural information through pre-trained encoders and overcomes the restricted relational reasoning of conventional attention via sparse expert activation. Our ablation studies show that while MoE within a single module yields marginal gains, cascading across multiple modules consistently improves performance, with a roughly linear relationship between configuration extremes and accuracy. We believe this experience is not only for AIDD but also for the broader field of AI4Science.

In virtual screening, HydrAffinity achieves enrichment comparable to or better than interaction-based methods in the top 5% of compounds, without requiring binding conformation preparation. This indicates that HydrAffinity is an effective pre-filter that substantially reduces the compound pool size while still retaining a large fraction of active compounds, thereby saving considerable time for subsequent precise molecular docking and interaction-based predictions, which are typically time-consuming. Furthermore, the lack of a universal winner across targets on LIT-PCBA suggests that no single molecular representation is optimal, implying that multi-modal fusion—rather than simple averaging—may improve generalization. Future work could focus on maximizing inter-modal complementarity. To specifically address the gap on extreme electrostatic properties, augmenting the training data with more extreme-charge ligands and exploring molecular representations or pre-trained models that with stronger charge-awareness could help preparation-free models better handle such edge cases.

In summary, this study clarifies the relative merits of molecular representation types, pre-training scale, and MoE design for PLA tasks. It provides systematic experimental evidence and a practical framework—HydrAffinity—for building more efficient and accurate preparation-free virtual screening systems. Beyond PLA prediction, the proposed evaluation framework and HydrAffinity architecture could be adapted to other biomolecular interaction tasks (e.g., protein– protein or nucleic acid–ligand binding), offering a generalizable pathway toward data-efficient and interpretable AI-driven discovery.

## Data and Software Availability

All protein–ligand complex structures and corresponding binding affinity data used in this study were sourced from the publicly available PDBbind database^34, 35^ (http://www.pdbbind.org.cn). The dataset is freely accessible to the research community without restrictions.

The molecular pretraining models employed in this work are publicly available. Specifically, the model weights can be accessed at the following repositories: ChemBERTa^9, 10^ (https://huggingface.co/DeepChem/models); PepDoRA^37^ (https://huggingface.co/ChatterjeeLab/PepDoRA); MolFormer^16^ (https://huggingface.co/ibm-research/MoLFormer-XL-both-10pct); MolAI^55^ (https://github.com/AnyoLabs/MolAI-Publication); SELFormer^56^ (https://huggingface.co/HUBioDataLab/SELFormer); UniMol^13^ (https://huggingface.co/dptech/Uni-Mol2); GeminiMol^17^ (https://huggingface.co/AlphaMWang/GeminiMol); ImageMol^57^ (https://github.com/ChengF-Lab/ImageMol); MaskMol^19^ (https://github.com/ZhixiangCheng/MaskMol). General pretrained image models are available through the torchvision and timm Python packages. Protein language models include: ESM-2^11^ (esm-fair Python package); ESM-3^12^ (https://huggingface.co/EvolutionaryScale/esm3-sm-open-v1); SaProt^30^ (https://huggingface.co/westlake-repl/SaProt_650M_AF2); ProSST-2048^14^ (https://huggingface.co/AI4Protein/ProSST-2048).

The DUDE-Z dataset can be found at https://dudez.docking.org. LIT-PCBA dataset can be reached at https://drugdesign.unistra.fr/LIT-PCBA.

Figure source data are provided with this paper. Assembled PDBBind Dataset and our HydrAffinity model weights can be found at https://zenodo.org/records/19336925.

## Code Availability

The code for reproducing the analyses and models is publicly available and has been deposited in GitHub at https://github.com/BHM-Bob/HydrAffinity_Review.

## Author Contributions

H.B. designed the study, developed the method, conducted the experiments and performed the analysis. H.B. and S.D. wrote the paper. All authors read and approved the manuscript.

## Conflicts of Interest

The authors declare no competing interests.

## Acknowledgements

This work was supported by the National Natural Science Foundation of China (32271307), the Natural Science Foundation of Gansu Province (26JRRA185, 26JRRA304).

## Supporting Information

Pretraining model data processing (Table S1), example of ligand molecular representations (Table S2), comprehensive summary of molecular representations (Table S3), Hydraffinity performance across datasets (Table S4), performance of Hydraffinity across different datasets under clean split (Table S5), training data and protocol comparison for methods evaluated on CASF-2016 (Table S6), UMAP of pretrained protein and ligand embeddings (Figure S1), SMILES embedding dimension vs. PLA performance (Figure S2), complementarity and conflict among receptor features (Figure S3), coupling between protein families and features (Figure S4), MoE predictor design (Figure S5), MoE ligand encoder design (Figure S6), MoE receptor encoder design (Figure S7), Hydraformer design (Figure S8), routing score variation vs. activated experts in MoE predictor (Figure S9), MoE activation patterns and performance (Figure S10), result in DUDE-Z and LIT-PCBA dataset (Figure S11), detailed information about ligand molecular representation pre-train model (Section S3), detailed mathematical formula description of MoE in this study (Section S4), performance on Davis, KIBA and BindingDB dataset (Section S16) (PDF).

## List of Abbreviations

CASF: Comparative Assessment of Scoring Functions
MLP: Multi-Layer Perceptron.
MoE: Mixture-of-Experts
PLA: Protein-ligand affinity.
RMSE: root-mean-square error.
SELFILES: SELF-referencIng Embedded Strings
SMILES: Simplified Molecular Input Line Entry System.

## Supplementary Information

### S1. Pretraining model data processing

**Table S1.**
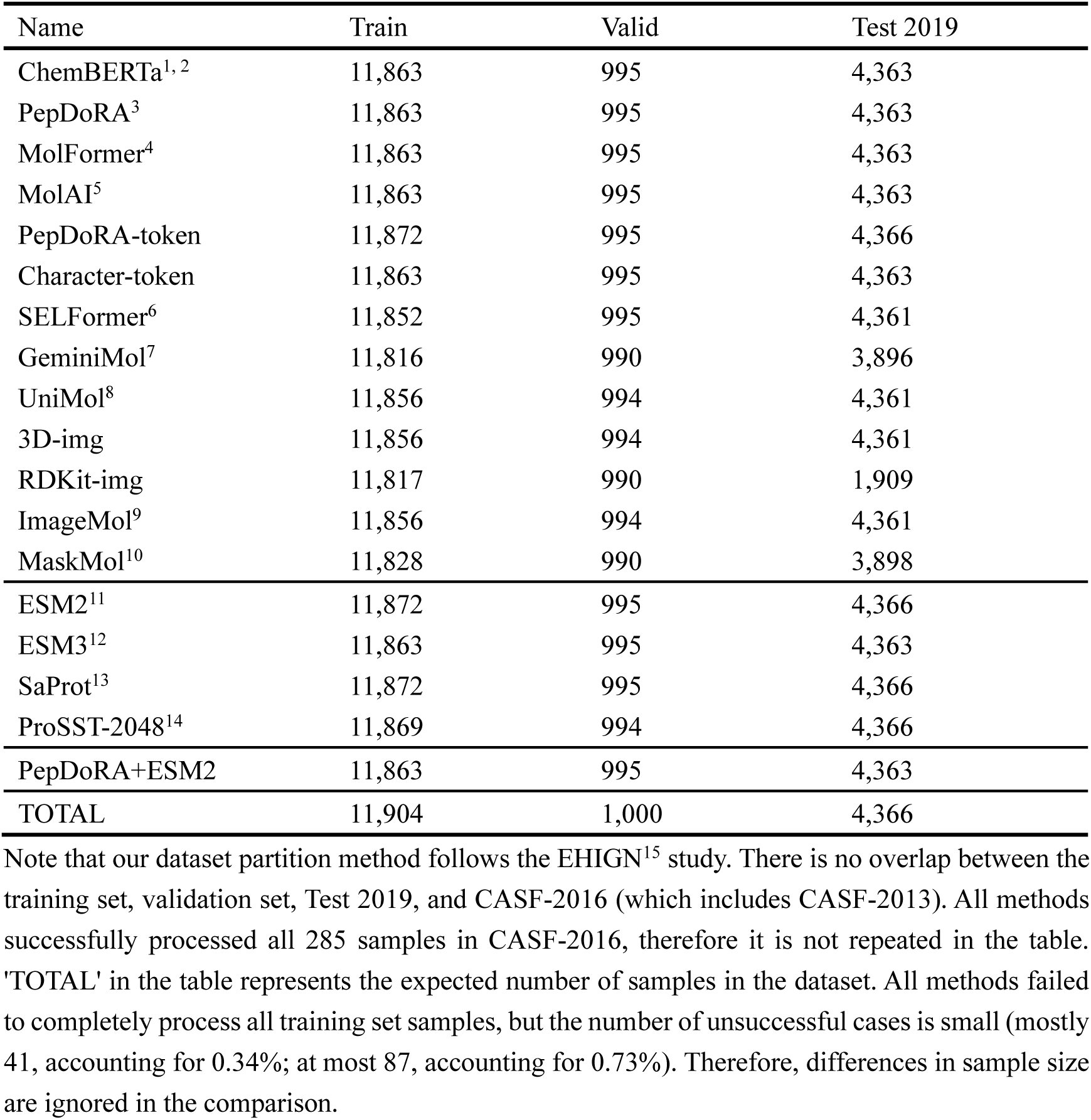
Data processing status of molecular representation pre-training models.

### S2. Example of ligand molecular representations

**Table S2.**
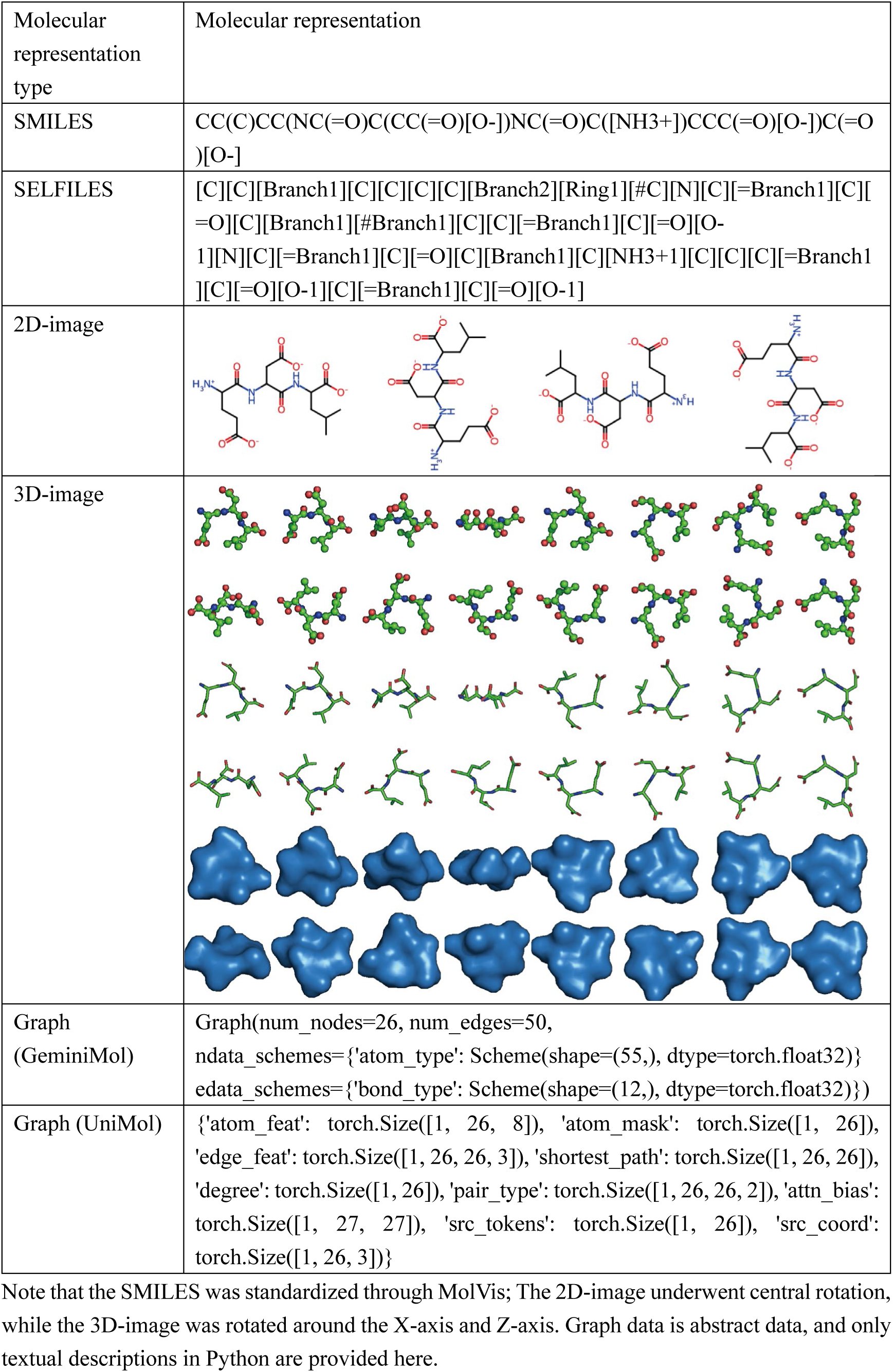
Ligand molecular representations of 1A30 in PDBbind v2016 dataset.

| Molecular representation type | Molecular representation |
| --- | --- |
| SMILES | CC(C)CC(NC(=O)C(CC(=O)[O-])NC(=O)C([NH3+])CCC(=O)[O-])C(=O)[O-] |
| SELFIES | [C][C][Branch1][C][C][C][C][Branch2][Ring1][#C][N][C][=Branch1][C][=O][C][Branch1][#Branch1][C][C][=Branch1][C][=O][O-1][N][C][=Branch1][C][=O][C][Branch1][C][NH3+1][C][C][C][=Branch1][C][=O][O-1][C][=Branch1][C][=O][O-1] |
| 2D-image |  |
| 3D-image |  |
| Graph (GeminiMol) | Graph(num_nodes=26, num_edges=50, ndata_schemes={'atom_type': Scheme(shape=(55,), dtype=torch.float32)} edata_schemes={'bond_type': Scheme(shape=(12,), dtype=torch.float32)}) |
| Graph (UniMol) | {'atom_feat': torch.Size([1, 26, 8]), 'atom_mask': torch.Size([1, 26]), 'edge_feat': torch.Size([1, 26, 26, 3]), 'shortest_path': torch.Size([1, 26, 26]), 'degree': torch.Size([1, 26]), 'pair_type': torch.Size([1, 26, 26, 2]), 'attn_bias': torch.Size([1, 27, 27]), 'src_tokens': torch.Size([1, 26]), 'src_coord': torch.Size([1, 26, 3])} |

Note that the SMILES was standardized through MolVis; The 2D-image underwent central rotation, while the 3D-image was rotated around the X-axis and Z-axis. Graph data is abstract data, and only textual descriptions in Python are provided here.

### S3. Detailed information about ligand molecular representation pre-train model

We utilized three types of molecular representations for ligands: linear text descriptors, molecular structures, and molecular images. For linear text descriptors, this study employed a series of large-scale pre-trained models based on SMILES: ChemBERTa^1, 2^ variants (10M-MLM, 77M-MLM, 77M-MTR, and 100M-MLM), PepDoRA^16^ (a fine-tuned model based on ChemBERTa-77M), MolFormer^4^ (1.3B model parameters), and MolAI^5^ (a pre-trained model based on an LSTM architecture). Additionally, we utilized SELFIES-based pre-trained models, such as SELFormer^6^. Furthermore, to explore the possibility of learning SMILES semantic spaces from scratch, we used PepDoRA’s tokenizer and a simple character-level SMILES vocabulary. For structural information of ligands, we employed UniMol v2^8^ (a graph neural network based on molecular conformations) and GeminiMol^7^ (a graph neural network for molecular representations trained using contrastive learning). For molecular image representation, we used ImageMol^9^ and MaskMol^10^, two image models pre-trained on RDKit-generated 2D molecular images (RDKit version 2024.9.5), with ImageMol based on a convolutional neural network architecture and MaskMol based on a transformer architecture. Additionally, we utilized general pre-trained CNNs^17^ and ViT^18^ models to process eight-view molecular images generated by PyMOL^19^ (version 3.0.0) in ball, stick, and surface modes, exploring the performance of image-based molecular representations as comprehensively as possible.

### S4. Detailed mathematical formula description of MoE in this study

#### Composition of MoE (Mixture-of-Experts)

a. Gate Network (Router): A linear layer

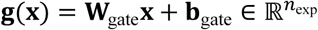

where **x** ∈ ℝ*^D^* is the hidden vector and *n*_exp_ is the number of experts.
b. Noisy Routing (Router Noise) (Used Only During Training):

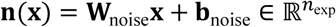 The output is transformed using softplus to obtain positive values, which are then multiplied by standard normal noise:

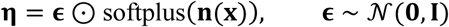 The final gating scores are **g**(**x**) + **η**.
c. Expert Networks: Subnetworks ℰ*_k_*(⋅), *k* = 1, …, *n*_exp_ that satisfy input/output dimension consistency. Gating scores are normalized under temperature *τ* > 0:

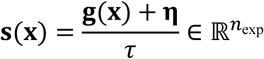 Temperature *τ* is a learnable parameter, ensuring positivity via *τ* = exp(log*τ*).

#### Top-k Selection

For each sample **x***_i_*, select the indices and scores of the top *k* = topk experts:

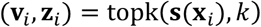

where **v***_i_* ∈ ℝ*^k^* are corresponding scores and **z***_i_* ∈ {1, …, *n*_exp_}*^k^* are expert indices.

During inference, gumbel-topk can be used to generate soft masks **m***_i_* ∈ [0,1]*^n^*^exp^.

#### Batch-forward Strategy

To reduce memory usage and achieve improved performance, MoE employs expert-grouped batch-forward.

Let the batch size be *N*, with each sample selecting *k* experts, resulting in *N* × *k* routes.

a. Expand Input

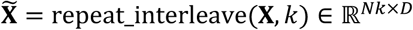
b. Aggregate by Expert For each unique expert *e*, take the corresponding input subset **X***e*, and compute expert outputs:

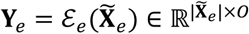

where *O* is the expert output dimension.
c. Reorganize Record the original batch position indices **idx***_e_*. Concatenate all **Y***_e_* and sort by **idx_e_**, obtaining:

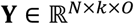

where **Y**[*i*, *j*,:] corresponds to the output of the j-th selected expert for sample *i*.

#### Gater Aggregation

a. sum:

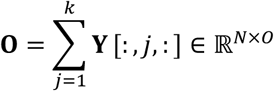
b. weighted First compute softmax over top-k scores:

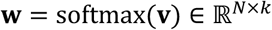

then perform weighted summation:

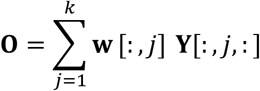

#### Load Balancing Loss

To prevent overuse of certain experts, we employ cross-entropy loss to constrain the gating distribution toward uniformity.

Let S ∈ ℝ*^N^*^×*n*exp^ be the gating scores (unnormalized), with target distribution 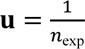. The cross-entropy is:

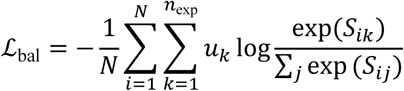

In implementation, this is equivalent to:

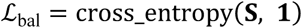

where all samples have target labels of 1, effectively enforcing uniform distribution constraints. The final training objective is:

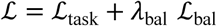

with *λ*_bal_ ∈ [10^−7^, 0.05] as a tunable loss scale.

#### Token-level MoE

Token-level MoE is implemented using the TokenMoE wrapper.

Given input **XX** ∈ ℝ*^N^*^×*L*×*D*^, first reshape to ℝ*^NL^*^×*D*^, pass through MoEPredictor, then reshape back:

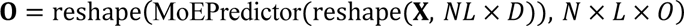

Here, each token independently routes to selected experts based on its hidden vector.

#### MLP Architecture

The MLP module is a standard two-layer MLP with LeakyReLU activations and dropout. Given an input tensor **X** ∈ ℝ^…×*D*^, the output O ∈ ℝ^…×*D*′^ is computed as:

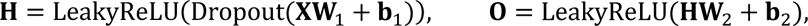

where W_1_ ∈ ℝ*^D^*^×^*^H^*, W_2_ ∈ ℝ*^H^*^×*D*′^, and the hidden dimension *H* defaults to 4*D*.

### S5. Comprehensive summary of molecular representations

**Table S3.**
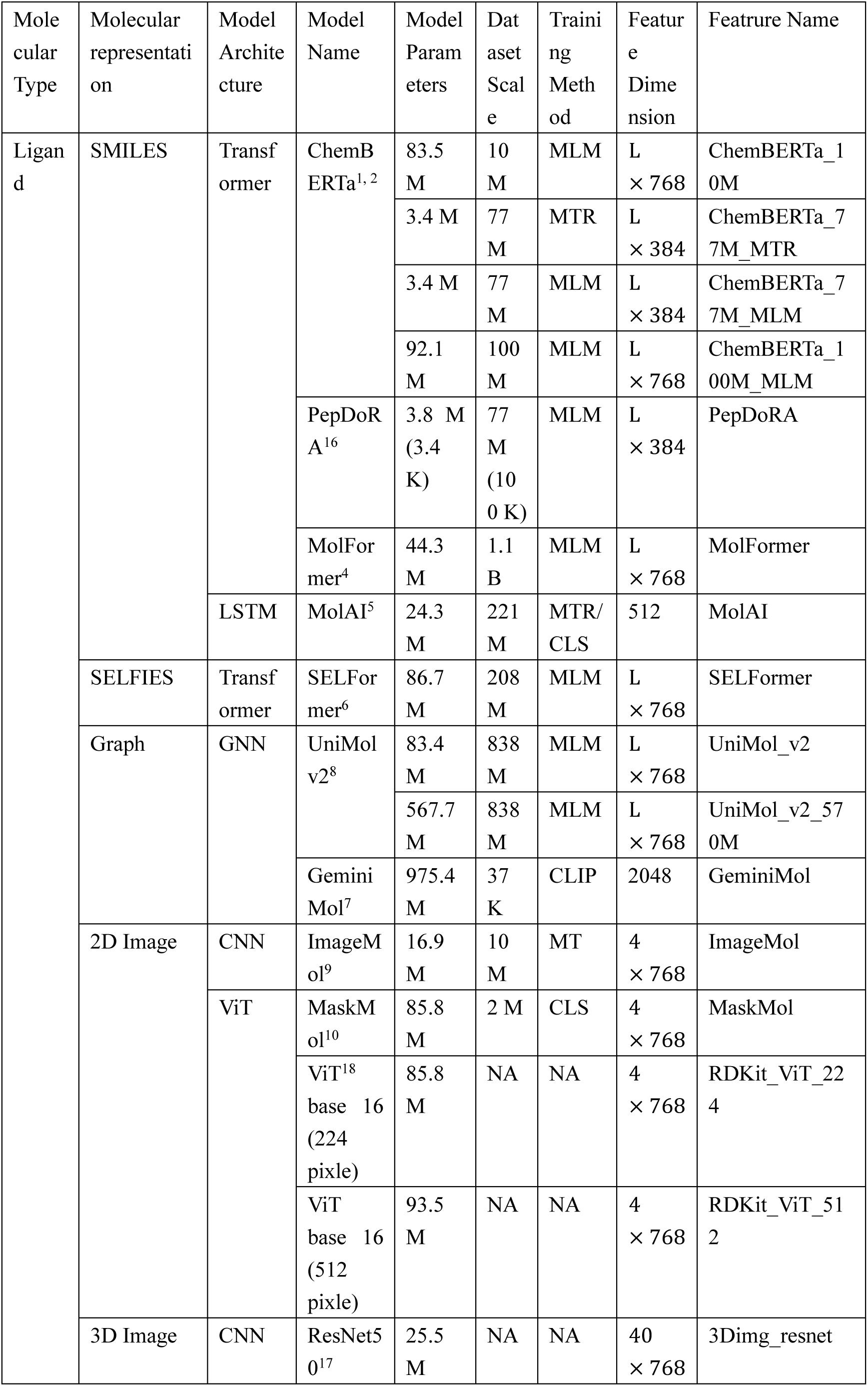
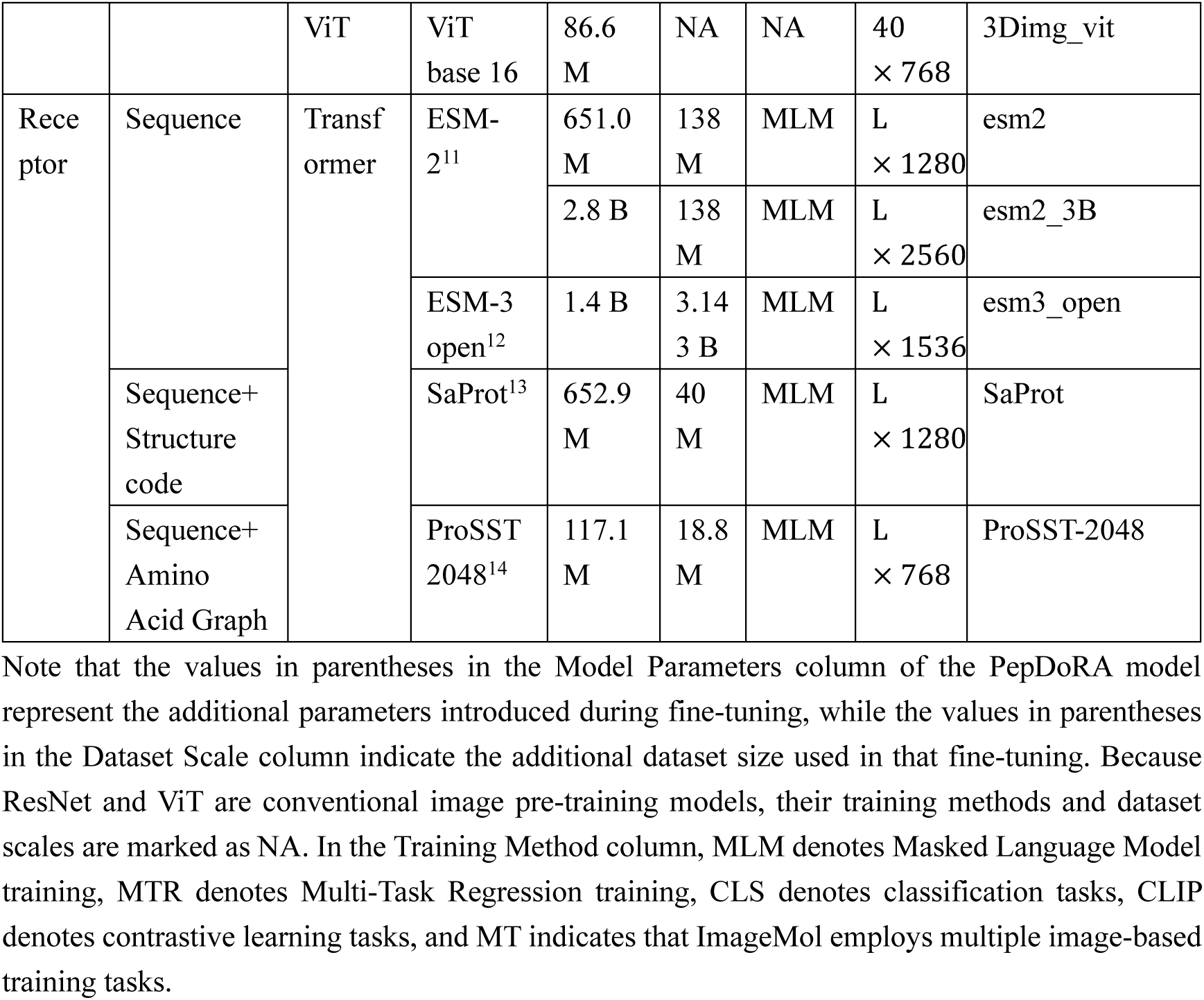
Molecular representation pretrain model summary.

### S6. UMAP of pretrained protein and ligand embeddings

**Figure S1:**
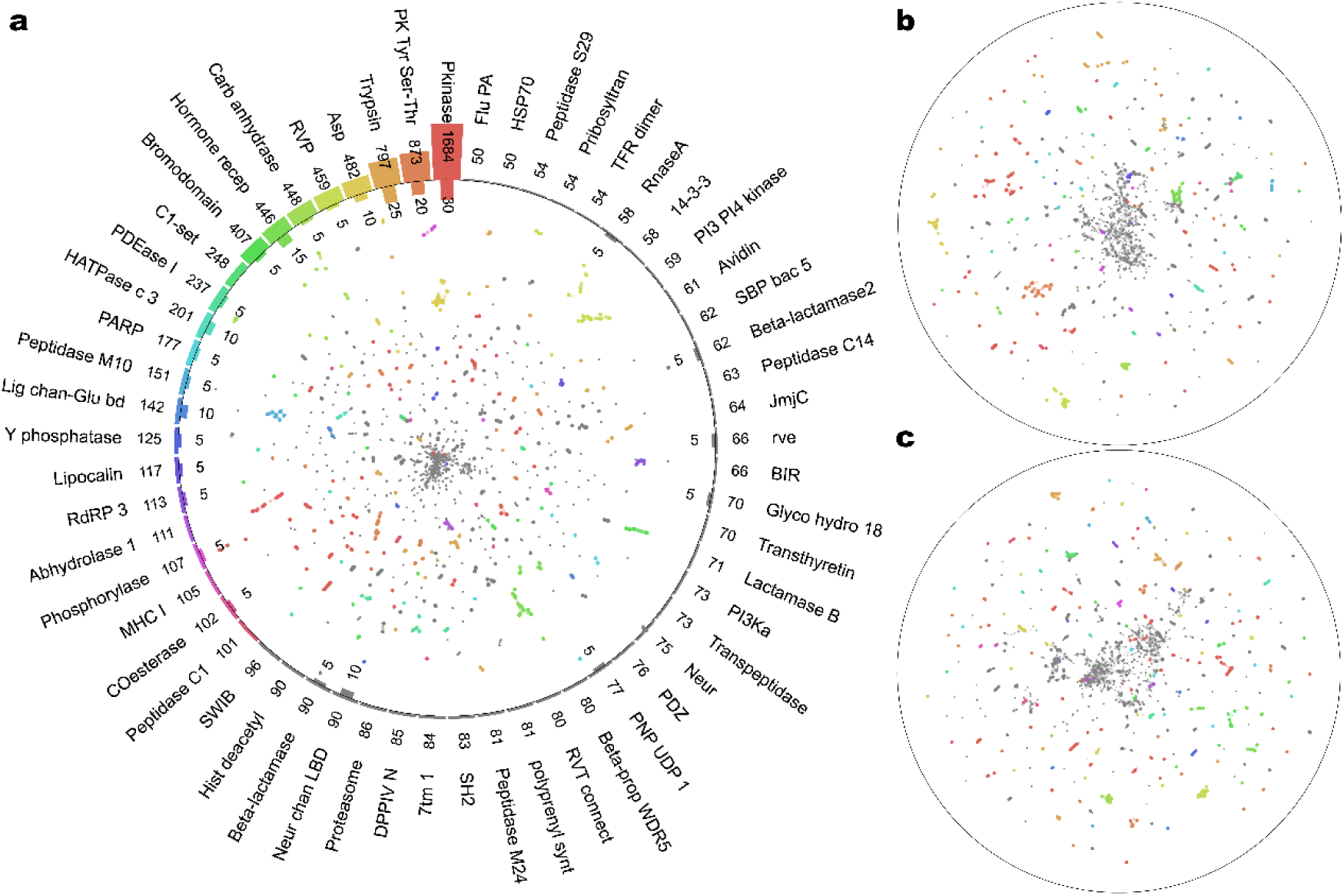
UMAP dimensionality-reduction result of protein embedding features. Proteins in the dataset were classified using the HMMER^20^ and Pfam databases^21^, and the high-dimensional embedding vectors generated by the pre-trained model (a: ESM2-3B^11^, b: ProSST-2048^14^, c: SaProt^13^) were subjected to UMAP for dimensionality reduction. (a) The scatter plot represents the 2-dimensional dimensionality reduction result of UMAP, with colors assigned only to the top 22 large protein families, and the rest shown in gray. Labels and bar charts are displayed only for families with 50 or more proteins. The bar charts and their numeric labels outside the circle represent the total number of proteins for each family in PDBbind v2019, while those inside the circle represent the number of proteins in CASF-2016^22^. The analysis demonstrates that, for large protein families, the pre-trained deep-learning model can effectively cluster proteins with similar amino-acid sequences into proximate groups within the high-dimensional space. This finding confirms that large-scale pre-training endows deep-learning models with the ability to recognize and characterize molecular similarity relationships, thereby providing robust theoretical support for the interaction-free PLA model.

### S7. SMILES embedding dimension vs. PLA performance

**Figure S2:**
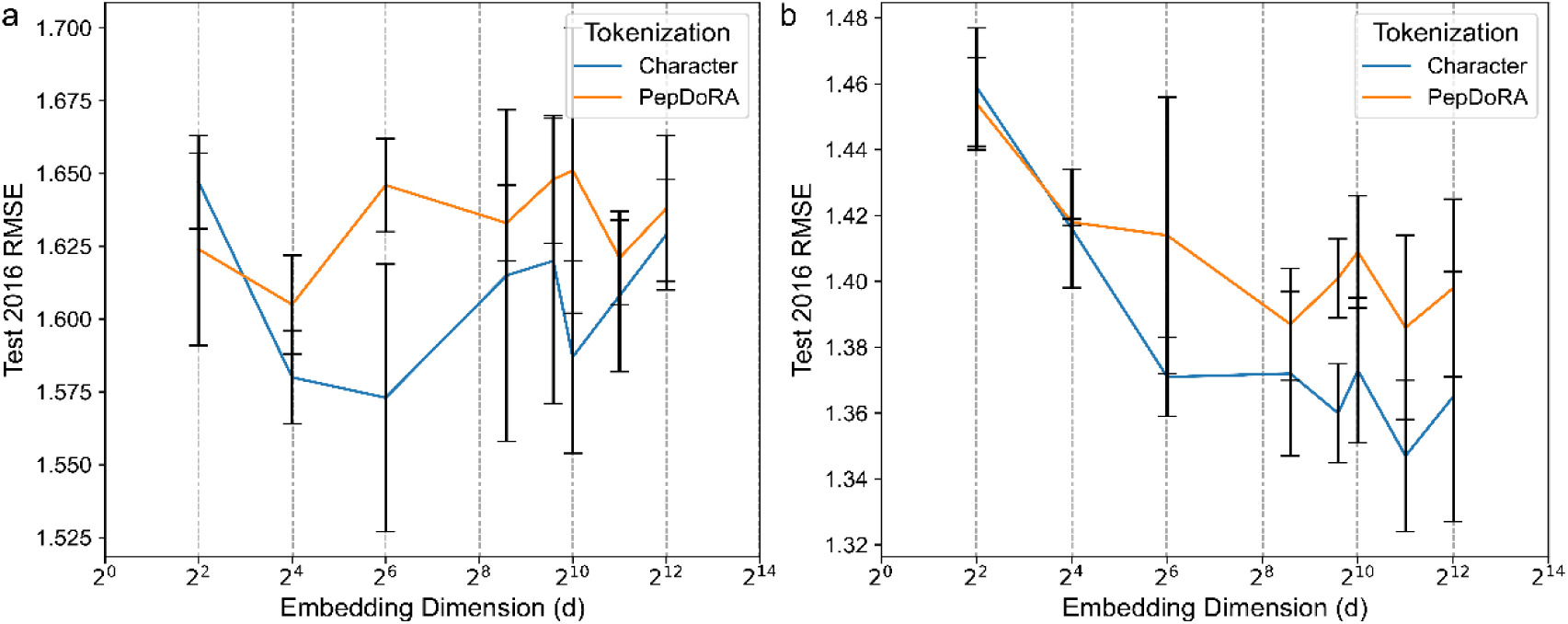
Relationship between the embedding dimension of SMILES token embeddings and PLA performance. (a) The relationship between the embedding dimension of SMILES token embeddings and PLA performance in single-modality settings. Overall, the performance of the custom character-level tokenizer outperforms that of PepDoRA’s tokenizer^16^; however, no significant trend emerges as the dimension changes. (b) The relationship between the embedding dimension of SMILES token embeddings and PLA performance in multi-modality settings. The receptor modality is ESM2-3B-33^11^. The overall performance differences are consistent with those in single-modality settings, but it can be observed that the performance of both tokenizers increases as the dimension grows, reaching a plateau around 2^8^ and showing no further significant improvements. Error bars in the figure represent standard deviation (std), with n=3.

### S8. Complementarity and conflict among receptor features

**Figure S3:**
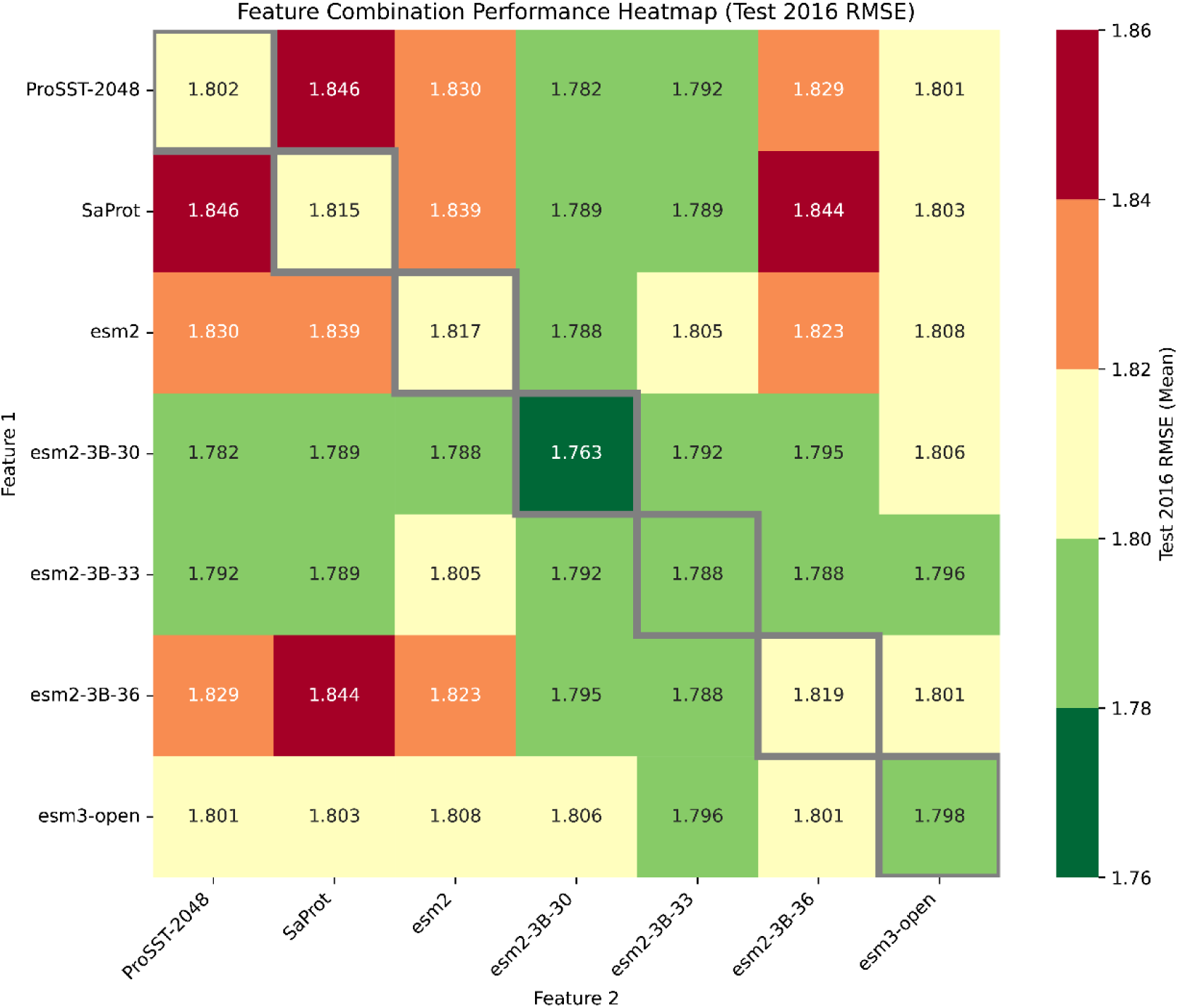
Individual and paired PLA performance of protein receptor features. Heat map of the RMSE test results on the CASF-2016 dataset was generated by training the receptor protein features either individually or in pairwise combinations. The results indicate that paired training often falls between the performance of the two individual features; however, many paired training cases perform worse than either individual feature alone, and no paired training case outperforms either one. This suggests a pronounced conflict between the receptor modalities. It is conceivable that dynamic adjustment of the weights or the direction of information flow between receptor modalities could optimize their complementarity, but a simple unified transformer architecture cannot achieve this.

### S9. Coupling between protein families and features

**Figure S4:**
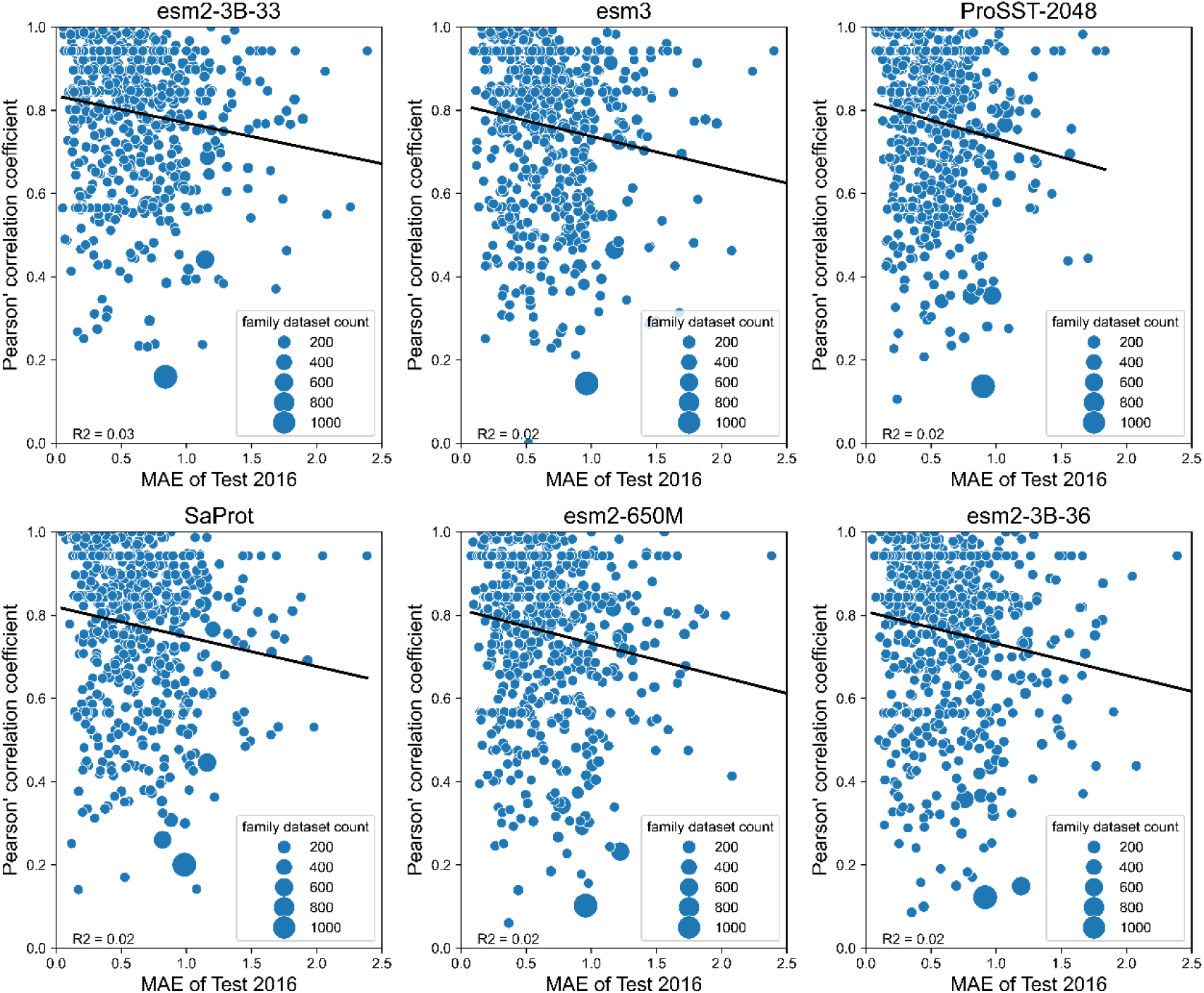
Relationship Between Receptor Pre-Training Embedding Similarity and Family Structure, and Its Impact on PLA Performance. Proteins from PDBbind v2019 were classified using the HMMER^20^ and pfam databases^21^. The TMalign algorithm^23^ was employed to calculate the TM-Score matrix for proteins within the same family. A similarity matrix based on pre-training embeddings was also constructed. The expanded similarities of these two matrices were computed, resulting in scatter plots correlating each family’s PLA task RMSE with their similarity scores. Notably, there is no significant relationship between the embedding-TM-Score correlation and PLA performance (*R*^2^ absolute value less than 0.05). This suggests that molecular insights learned by pre-training models may not directly translate into superior PLA performance.

### S10. MoE predictor design

**Figure S5:**
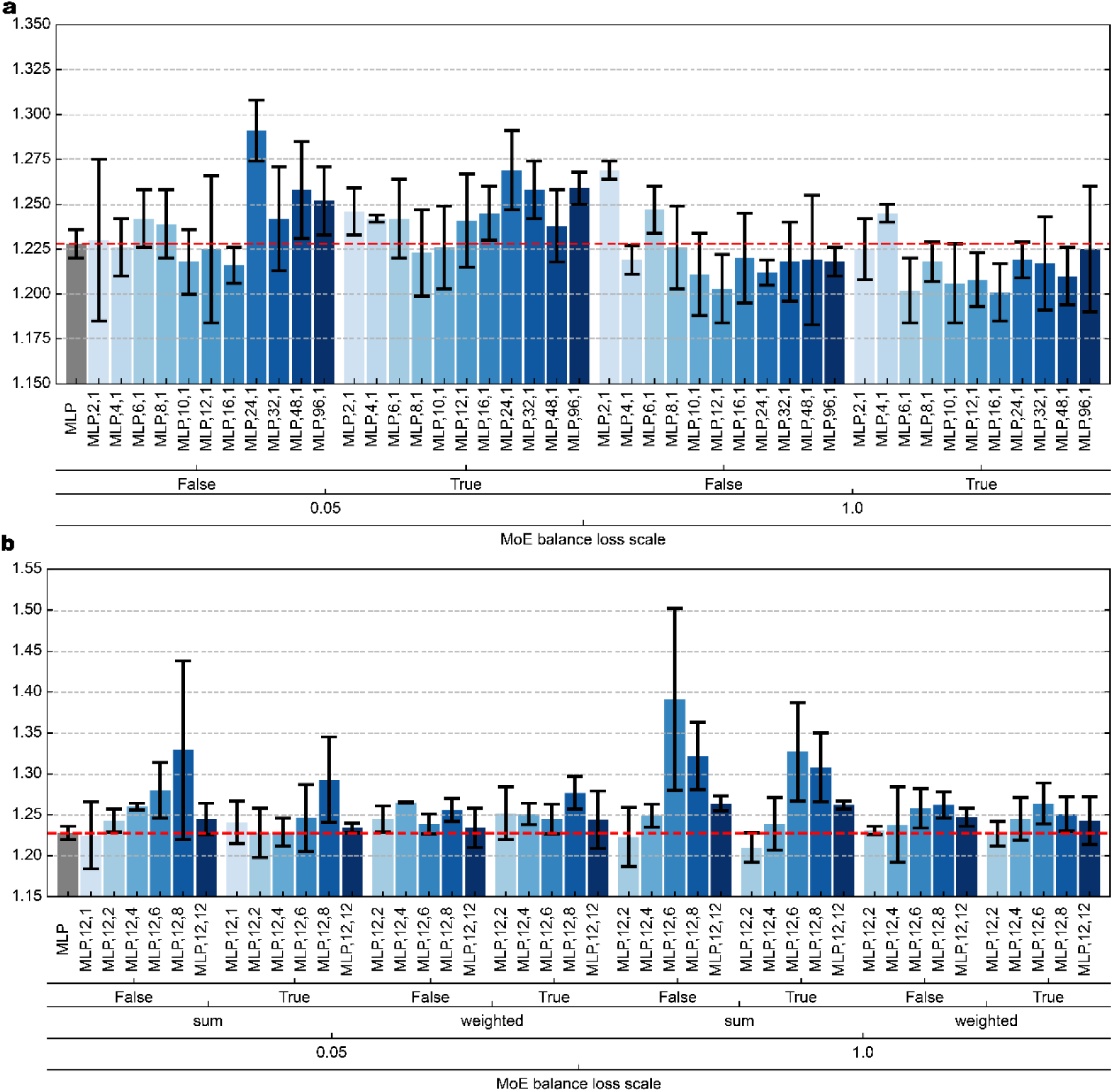
Performance of the MoE predictor on CASF-2016. The x-axis labels, MLP, M, N, represent a total of M experts with N active experts; False and True indicate whether router noise is enabled or disabled; sum and weighted denote the router aggregation strategies; 0.05 and 1.0 correspond to the MoE balance loss scale. (a) Performance of the CASF-2016 model (RMSE) under different total expert numbers when the number of activated experts in the predictor is fixed at 1. PepDoRA and esm2-3B-33 were used as ligand and receptor modalities respectively, the model’s CASF-2016 performance first improves and then deteriorates as the total number of experts increases from 2 to 96. Additionally, increasing the scaling factor of the load balancing loss from 0.05 to 1 leads to a more stable and effective improvement in model performance. This suggests that load balancing plays a significant role in enhancing the performance of MoE models. (b) Performance of the CASF-2016 model (RMSE) under different numbers of activated experts when the total number of experts in the predictor is fixed at 12. We observe that under the arithmetic summation aggregation strategy, model performance gradually decreases as the number of activated experts increases from 1 to 12; however, under the weighted summation aggregation strategy, performance remains largely unchanged. This may indicate that as the number of activated experts increases, there is an activation of unnecessary experts, and a simple weighted summation aggregation strategy fails to effectively filter out these “harmful” outputs. The weighted summation approach can mitigate this issue by assigning lower weights to less important or even detrimental expert activations. Error bars in the figure represent standard deviation (std), with n=3.

### S11. MoE ligand encoder design

**Figure S6:**
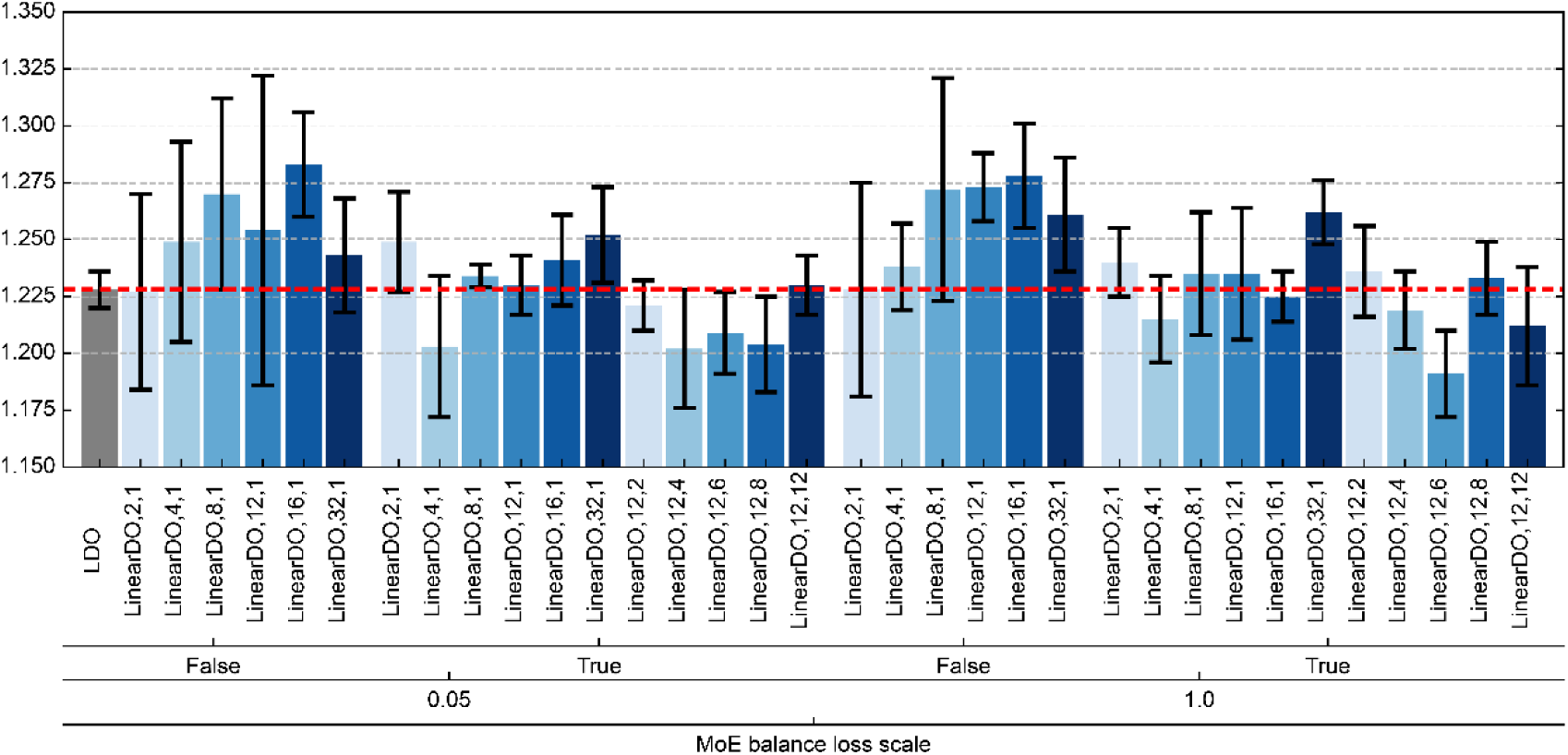
Performance of the MoE ligand encoder under different configurations. The x-axis labels, LinearDO, M, N, represent a total of M experts with N active experts; False and True indicate whether token-level MoE router is enabled or disabled; 0.05 and 1.0 denote the MoE balance loss scale. PepDoRA and esm2-3B-33 were used as the modalities for the ligand and receptor, respectively, with only the ligand encoder employing a MoE structure, where the router employs a sum aggregation strategy. Four variables were systematically investigated: (1) the scaling factor of the load balancing loss in the ligand MoE encoder (0.05 and 1), (2) whether token-level MoE was enabled (yes or no), (3) the total number of experts in the ligand MoE encoder (2, 4, 8, 12, 16, and 32), and (4) the number of activated experts in the ligand MoE encoder (with a fixed total expert count of 12; activated counts: 1, 2, 4, 6, 8, and 12). Systematic experiments revealed that token-level MoE routing strategies generally outperformed sequence-level MoE routing strategies. Additionally, an increasing number of MoE experts overall correlated with poorer model performance. Regarding the scaling factor of the load balancing loss in the MoE encoder, no significant differences were observed across configurations, suggesting that developers may not need to overly focus on this hyperparameter. Error bars in the figure represent standard deviation (std), with n=3.

### S12. MoE receptor encoder design

**Figure S7:**
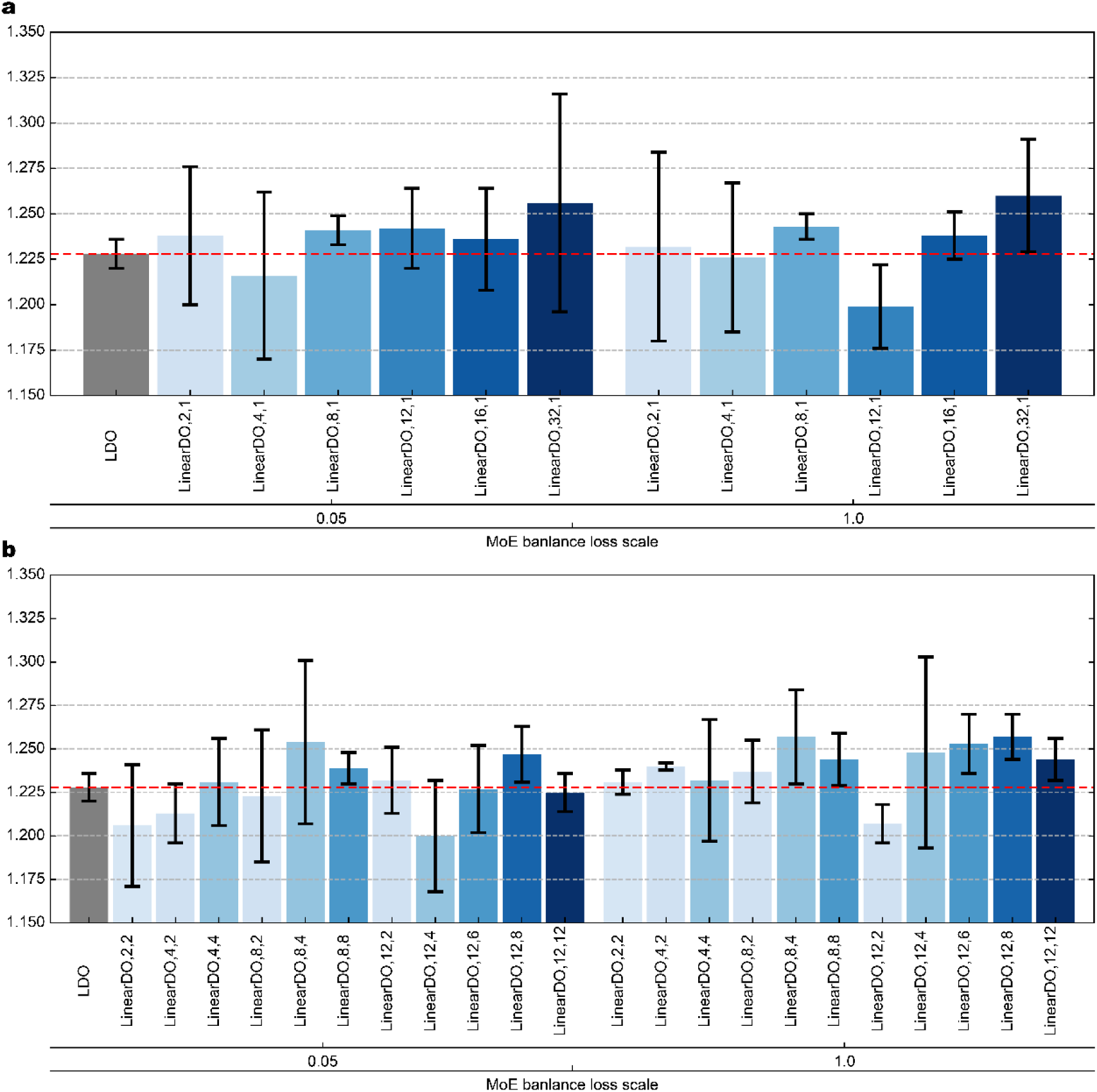
MoE receptor encoder. The x-axis labels, LinearDO, M, N, represent a total of M experts with N active experts; 0.05 and 1.0 denote the MoE balance loss scale. PepDoRA and esm2-3B-33 were used as ligand and receptor modalities, respectively, with only the receptor encoder employing the MoE structure. (a) Two variables were systematically investigated: (i) the scaling factor of the load balance loss for the receptor MoE encoder (0.05 and 1), and (ii) the total number of experts in the receptor MoE encoder (2, 4, 8, 12, 16, and 32). Systematic experiments revealed that unlike the ligand MoE encoder, variations in the total number of experts did not significantly impact or cause fluctuations in model performance. Similarly, the scaling factor for the load balance loss in the receptor MoE encoder showed no overall differences between 0.05 and 1. (b) Two variables were systematically investigated: (i) the scaling factor of the load balance loss for the receptor MoE encoder (0.05 and 1), and (ii) the number of active experts and total number of experts in the receptor MoE encoder (2, 4, 8, and 12). Systematic experiments showed that when the total number of experts is small or the activation ratio is less than 50%, the model tends to achieve better performance. Additionally, the load balance loss scaling factor did not result in overall differences in this case either. Error bars in the figure represent standard deviation (std), with n=3.

### S13. Hydraformer design

**Figure S8:**
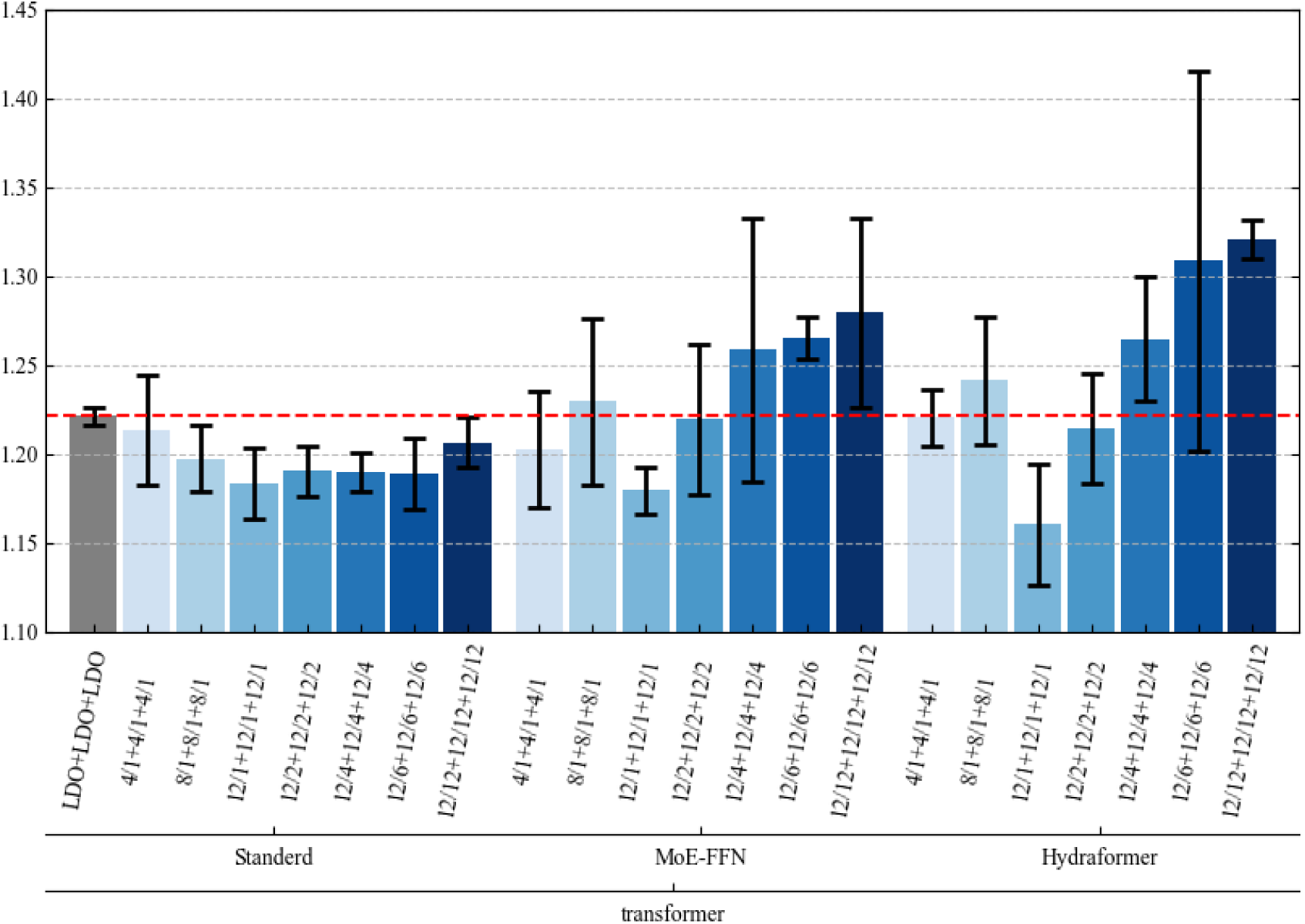
Performance Analysis of Different Transformers Under MoE Architecture. The x-axis labels, M/N + M/N + M/N, represent a total of M experts with N active experts for ligand MoE encoder, receptor MoE encoder and MoE predictor; Standerd, MoE − FFN and Hydraformer donate the transformer type.

### S14. Hydraffinity performance across datasets

**Table S4.** Performance of Hydraffinity on each metrics.

| Name | RMSE | MAE | R<sub>p</sub> | CI |
| --- | --- | --- | --- | --- |
| CASF-2013 |  |  |  |  |
| Bimodal | 1.359 (0.030) | 1.041 (0.028) | 0.812 (0.010) | 0.811 (0.006) |
| Multimodal | 1.331 (0.047) | 1.047 (0.043) | 0.824 (0.017) | 0.814 (0.012) |
| CASF-2016 |  |  |  |  |
| Bimodal | 1.161 (0.034) | 0.885 (0.030) | 0.846 (0.009) | 0.826 (0.006) |
| Multimodal | 1.170 (0.035) | 0.905 (0.018) | 0.844 (0.011) | 0.824 (0.004) |
| PDBbind v2019 houdout |  |  |  |  |
| Bimodal | 1.454 (0.020) | 1.135 (0.011) | 0.642 (0.007) | 0.725 (0.003) |
| Multimodal | 1.493 (0.031) | 1.169 (0.025) | 0.623 (0.010) | 0.719 (0.004) |

Note that the Bimodal variant uses PepDoRA as the ligand modality and esm2-3B-33 as the receptor modality. The Multimodal variant uses PepDoRA as the ligand modality, and esm2-3B-33, SaProt, and ProSST-2048 as the receptor modalities, employing a Mixture of Experts (MoE) configuration (the predictor, ligand encoder, and receptor encoder are all MoEs with a total of 12 experts and 1 activated expert; the ligand encoder MoE uses a token-level routing strategy, and the transformer is a three-modal-path Hydraformer).

**Table S5.** Performance of Hydraffinity across different datasets under clean split.

| Name | CASF-2013 |  | CASF-2016 |  |
| --- | --- | --- | --- | --- |
|  | RMSE | R<sub>p</sub> | RMSE | R<sub>p</sub> |
| GEMS | - | - | **1.308 (0.026)** | **0.803 (0.006)** |
| GenScore | - | - | 1.362 (0.029) | 0.780 (0.015) |
| Pafnucy | - | - | 1.484 (0.038) | 0.746 (0.015) |
| HydrAffinity (Bimodal) | 1.575 (0.067) | 0.735 (0.026) | 1.416 (0.038) | 0.760 (0.015) |
| HydrAffinity (Multimodal) | **1.564 (0.021)** | **0.743 (0.009)** | **1.385 (0.049)** | **0.772 (0.020)** |

Note that the data for GEMS<sup>24</sup>, GenScore<sup>25</sup>, and Pafnucy<sup>26</sup> are sourced from the GEMS publication. GEMS and GenScore are GNN-based interaction-based methods, while Pafnucy is a 3D CNN-based interaction-based method. In HydrAffinity, the Bimodal variant uses PepDoRA as the ligand modality and esm2-3B-33 as the receptor modality, with the architecture consistent with the state-of-the-art model described in the main text. The Multimodal variant uses PepDoRA as the ligand modality, and esm2-3B-33, SaProt, and ProSST-2048 as the receptor modalities, employing a Mixture of Experts (MoE) configuration (the predictor, ligand encoder, and receptor encoder are all MoEs with a total of 12 experts and 1 activated expert; the ligand encoder MoE uses a token-level routing strategy, and the transformer is a three-modal-path Hydraformer). Performance data are presented as mean (std), n=5. Bold text indicates the best performance.

### S15. Training data and protocol comparison for methods evaluated on CASF-2016

In Table 1, all compared methods train on the PDBbind series (primarily v2016) and evaluate on the CASF-2016 core set. The core de-duplication principle — removing samples that overlap with the test set — is consistently applied. Variations in reported training-set size arise mainly from differences in the PDBbind version used and, more importantly, from inevitable sample losses during data encoding. For instance, graph-based methods may discard complexes that fail RDKit sanitization, and Dynaformer further filters out complexes unsuitable for molecular dynamics simulation. These encoding-dependent losses are inherent to each method and do not reflect differences in the train/test split design. An intuitive and detailed comparison is provided in Table S6.

**Table S6.**
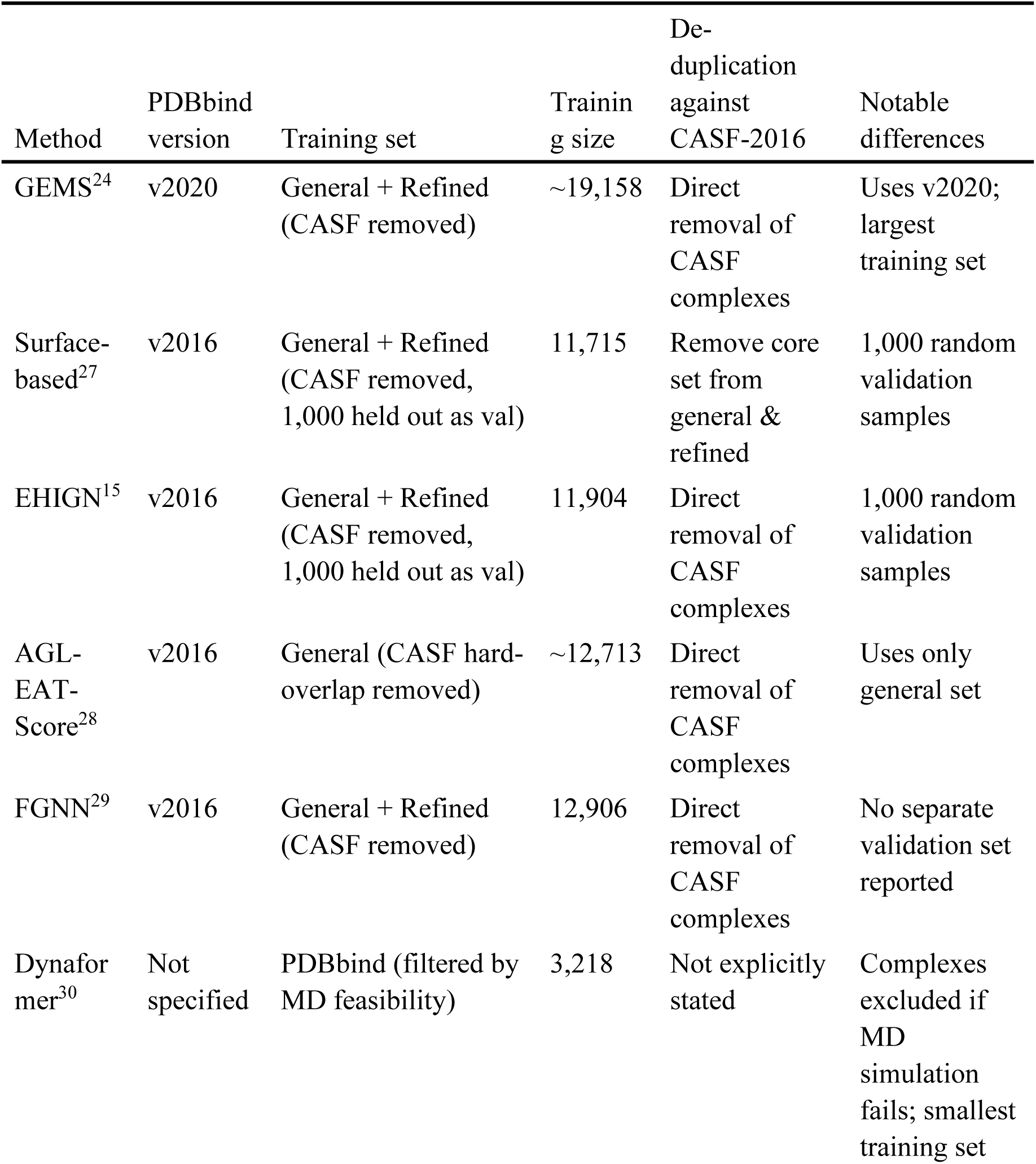
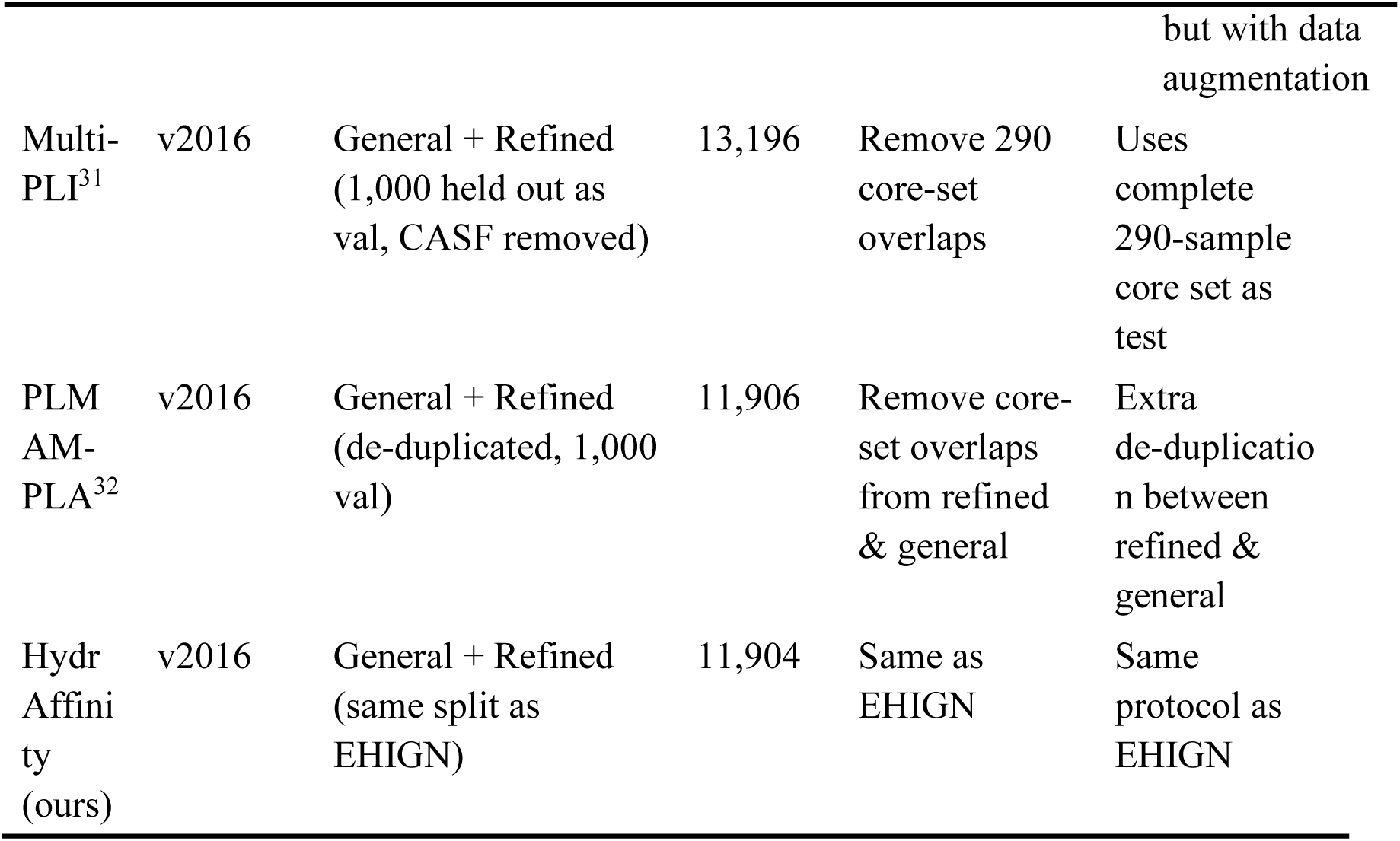
Comparison of dataset for methods in Table 1.

### S16. Performance on Davis, KIBA and BindingDB dataset

To examine the generalizability of the findings regarding pretrained model evaluation and MoE design, we additionally evaluated our model on the Davis^33^, KIBA^34^, and BindingDB^35^ datasets, which are widely used by many interaction-free models^36–38^.

Because these three datasets contain relatively few proteins and ligands, we used the best-performing ligand (GeminiMol) and receptor (esm3-3B-30) pretrained models for feature extraction, but replaced the complex modality encoders with a single linear projection layer that maps the pretrained features to the unified hidden dimension. We also replaced the Hydraformer with a standard Transformer encoder and retained only the MoE predictor (8 MLP experts with routing noise and top-1 gating) to mitigate overfitting. To improve ranking power on these small datasets, we additionally introduced a pairwise ranking loss with a scale of 2. All other training details are the same as those used for the PDBbind dataset.

Using the performance data collected by DeepDTAGen^36^, we found that our model achieves the best MSE among all compared methods, and the best CI on the Davis and KIBA datasets, while it does not show an advantage in *r*^2^ and AUPR (Table S7). This indicates that the pretrained model selection results and the design principles of MoE identified in this study can be readily transferred to other datasets with satisfactory performance.

**Table S7.** Performance on Davis, KIBA and BindingDB dataset.

| Dataset | Method | CI (std) | MSE (std) | $r_m^2$ (std) | AUPR (std) |
| --- | --- | --- | --- | --- | --- |
| KIBA | DeepDTA<sup>39</sup> | 0.863 (0.002) | 0.194 | 0.630 (0.009) | 0.788 (0.004) |
|  | GDilatedDTA<sup>40</sup> | 0.876 | 0.156 | **0.775** | - |
|  | DeepDTAGen<sup>41</sup> | **0.897 (0.003)** | 0.146 (0.008) | 0.765 (0.005) | **0.843 (0.001)** |
|  | **HydrAffinity (L)** | 0.896 (0.0019) | **0.140 (0.0013)** | 0.729 (0.0132) | 0.657 (0.0046) |
| Davis | DeepDTA<sup>39</sup> | 0.878 (0.004) | 0.261 | 0.630 (0.017) | 0.714 (0.010) |
|  | GDilatedDTA<sup>40</sup> | 0.885 | 0.237 | 0.686 | - |
|  | DeepDTAGen<sup>41</sup> | 0.890 (0.004) | 0.214 (0.006) | **0.705 (0.001)** | **0.772 (0.002)** |
|  | **HydrAffinity (L)** | **0.898 (0.0016)** | **0.203 (0.0015)** | 0.687 (0.0156) | 0.535 (0.0056) |
| BindingDB | DeepDTA<sup>39</sup> | 0.844 (0.003) | 0.633 (0.0002) | 0.633 (0.004) | 0.710 (0.0001) |
|  | GDilatedDTA<sup>40</sup> | 0.868 (0.001) | 0.483 (0.002) | 0.730 (0.001) | 0.820 (0.00021) |
|  | DeepDTAGen<sup>41</sup> | 0.876 (0.004) | 0.458 (0.002) | **0.760 (0.003)** | **0.870 (0.004)** |
|  | **HydrAffinity (L)** | **0.883 (0.0006)** | **0.430 (0.0060)** | 0.715 (0.0095) | 0.696 (0.0020) |

Note: Performance data for DeepDTA, GDilatedDTA and DeepDTAGen are all taken from the published work of DeepDTAGen. HydrAffinity (L) denotes the lightweight configuration described above (standard Transformer backbone + MoE predictor with 8 experts and top-1 gating).

### S17. Routing score variation vs. activated experts in MoE predictor

**Figure S9:**
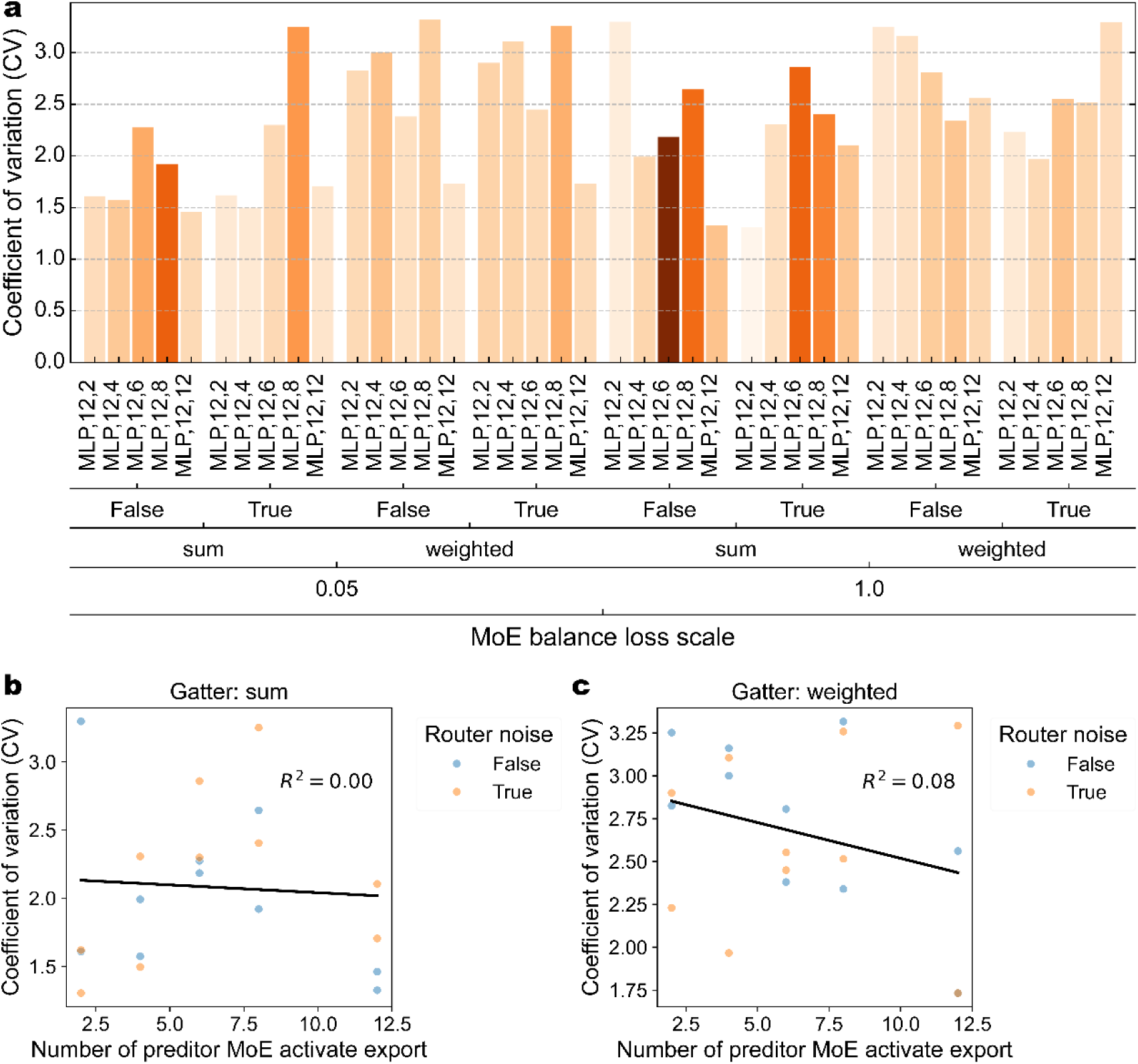
Number of Activated Experts and Coefficient of Variation of Activation Scores in MoE Routing. Relationship between the number of activated experts and the coefficient of variation of routing scores when the MoE predictor uses a fixed total of 12 experts. PepDoRA and esm2-3B-33 were used as the ligand and receptor modalities, respectively, with only the predictor employing a MoE architecture. Three variables were set: the scaling factor for the load balancing loss of the receptor MoE encoder (0.05 and 1), the routing aggregation strategy (arithmetic summation or weighted summation), and the number of activated experts (2, 4, 6, 8, and 12). (a) Experimental configuration and bar chart of the routing coefficient of variation. Color represents the relative value of CASF-2016 RMSE. Scatter plots showing the relationship between the number of activated experts and the coefficient of variation of routing scores when using arithmetic summation (b) and weighted summation (c) aggregation strategies. Systematic experiments reveal that, regardless of whether the arithmetic summation or weighted summation strategy is used, the number of activated experts is independent of the coefficient of variation of the routing scores.

### S18. MoE activation patterns and performance

**Figure S10:**
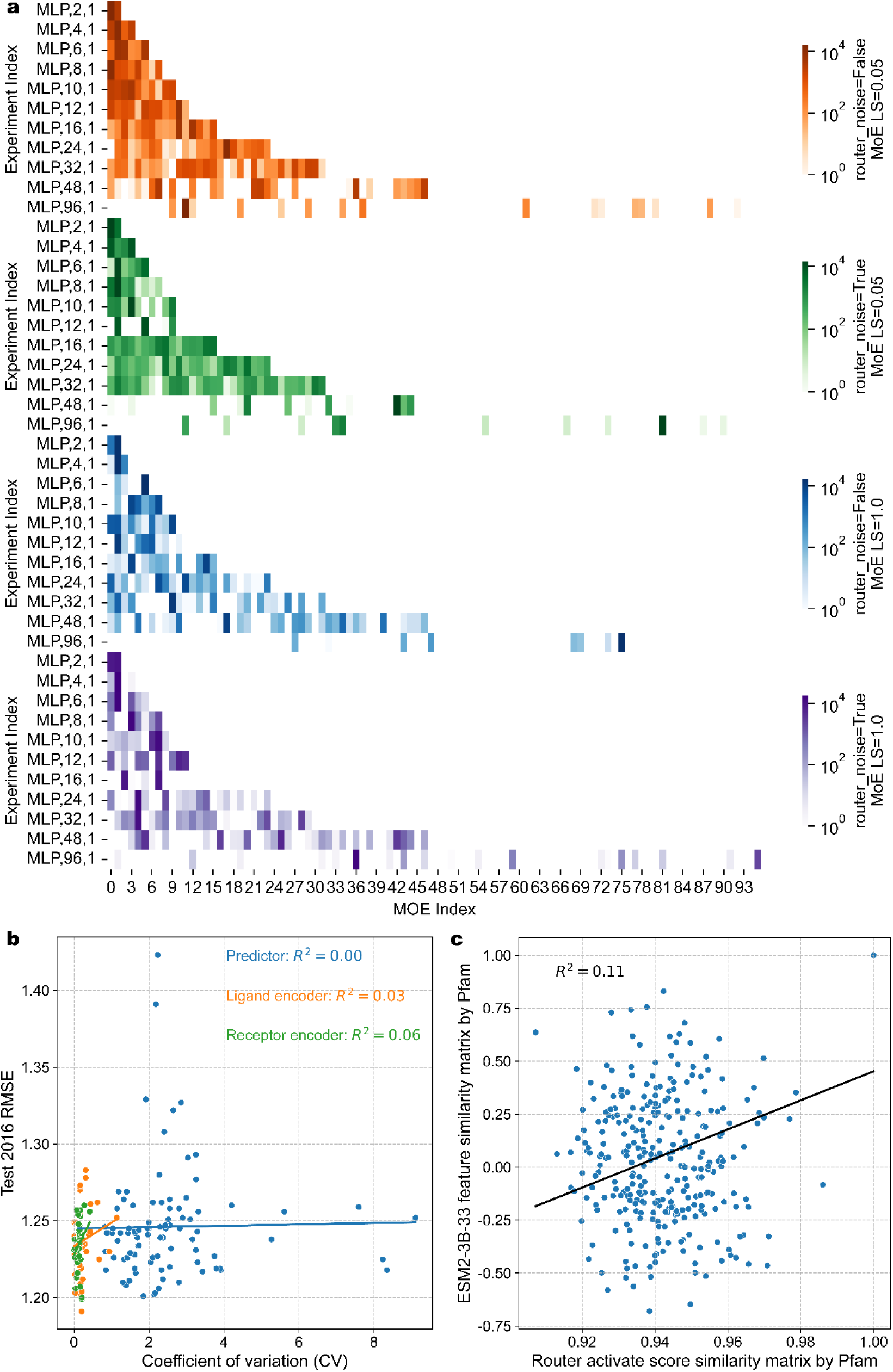
Relationship between MoE activation and performance. PepDoRA and esm2-3B-33 were used as the ligand and receptor modalities. a, Heatmap of expert activation patterns in the Predictor MoE. This panel counts the activation frequency of expert modules across all datasets. It can be observed that as the total number of experts increases, the MoE does not adequately activate and utilize all experts. Particularly when the total number of experts is 48 or more, only about half or even a small number of experts are activated. b, Relationship between the coefficient of variation (CV) of MoE router activation scores and model performance (RMSE) on CASF-2016. It can be seen that there is no correlation between the CV of activation scores and model performance for the predictor, ligand modality encoder, or receptor modality encoder. c, Proteins in the dataset were classified into various protein families using HMMER^20^ and Pfam^21^. This experiment is the same as Figure S4c in the main text. This experiment uses a Receptor modality encoder with an MoE architecture only (total experts: 12, activated experts: 4, using router noise, using the sum routing aggregation strategy, Test 2016 performance: 1.200 (0.032), mean (std)). The feature family similarity matrix was calculated using embeddings from esm2-3B-33; the router family similarity matrix was also calculated using router activation scores; a scatter plot of the flattened vectors from these two matrices was drawn and the Pearson correlation coefficient was calculated. It can be found that there is no correlation between the two (*R*^2^ = 0.11).

### S19. Result in DUDE-Z and LIT-PCBA dataset

**Figure S11:**
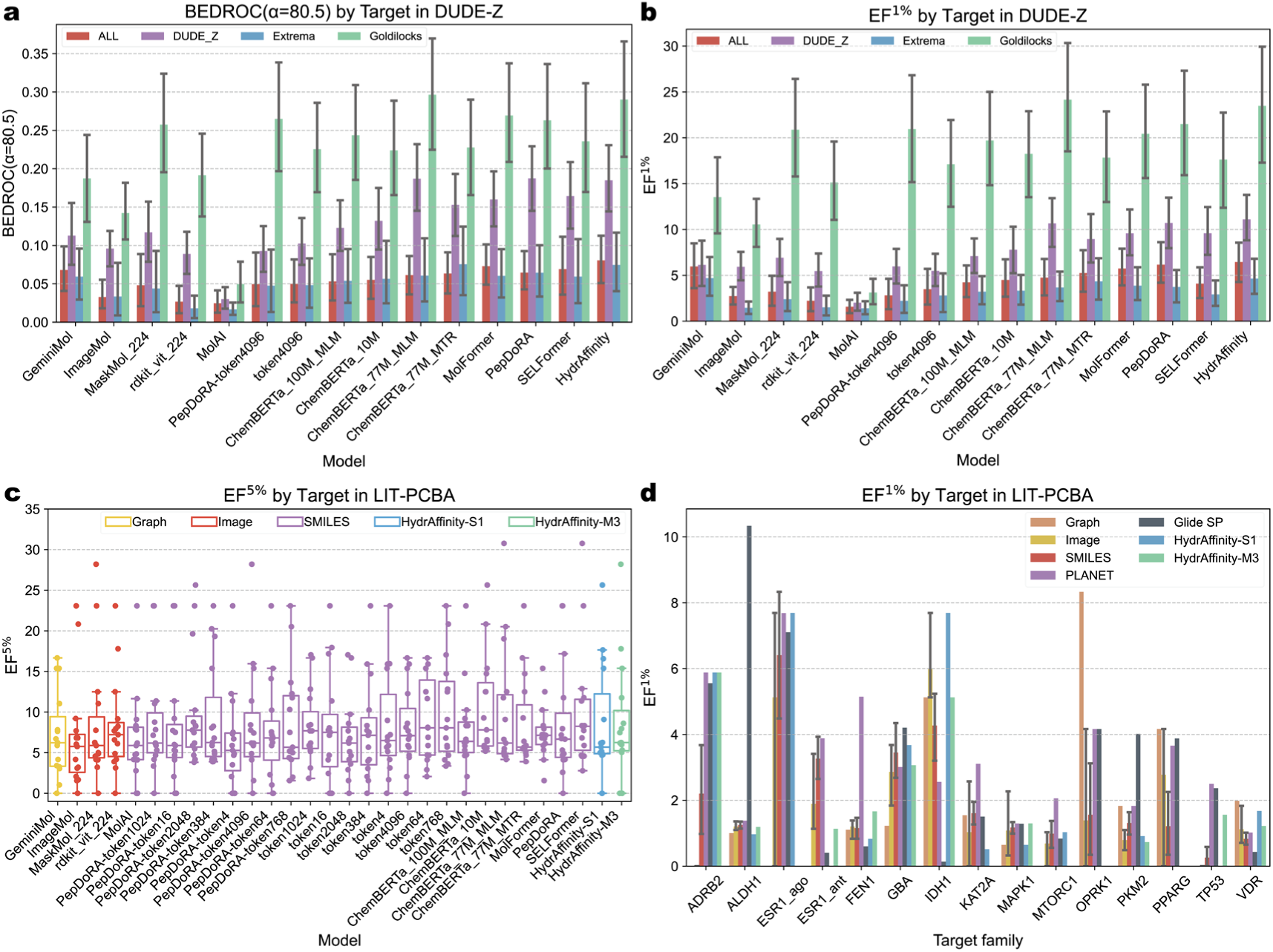
Result in DUDE-Z and LIT-PCBA dataset. a (BEDROC (*α* = 80.5)), b (EF^1%^) results in DUDE-Z dataset. It can be found that the performance of MoAI is the lowest under both metrics, and all pre-trained models perform best on Goldilocks. c, EF^5%^ results in LIT-PCBA by each ligand modal. Unlike DUDE-Z, MolAI did not show a significant disadvantage on the LIT-PCBA. d, EF^1%^ results in LIT-PCBA by each ligand molecular representation and method. HydrAffinity-S1 and HydrAffinity-M3 stands for bimodal architecture and multimodal architecture in Table 2 of main text. Performance of PLANET and Glide SP are taken from PLANET works. Error bars on the bar charts represent 95% confidence level.

## Notes

https://zenodo.org/records/20595631

### Competing Interest Statement

The authors have declared no competing interest.

## References

(1) Warr, W. A.; Nicklaus, M. C.; Nicolaou, C. A.; Rarey, M. Exploration of Ultralarge Compound Collections for Drug Discovery. Journal of Chemical Information and Modeling 2022, 62 (9), 2021–2034. DOI: 10.1021/acs.jcim.2c00224.

(2) Wang, D. D.; Wu, W.; Wang, R. Structure-based, deep-learning models for protein-ligand binding affinity prediction. Journal of Cheminformatics 2024, 16 (1), 2. DOI: 10.1186/s13321-023-00795-9.

(3) Yang, Z.; Zhong, W.; Lv, Q.; Dong, T.; Chen, G.; Chen, C. Y.-C. Interaction-Based Inductive Bias in Graph Neural Networks: Enhancing Protein-Ligand Binding Affinity Predictions From 3D Structures. IEEE Transactions on Pattern Analysis and Machine Intelligence 2024, 1–18. DOI: 10.1109/TPAMI.2024.3400515.

(4) Cui, H.; Su, Y.; Dean, T. J.; Yu, T.; Zhang, Z.; Peng, J.; Shukla, D.; Zhao, H. Enzyme specificity prediction using cross attention graph neural networks. Nature 2025. DOI: 10.1038/s41586-025-09697-2.

(5) Tao, H.; Wang, X.; Huang, S.-Y. An interaction-derived graph learning framework for scoring protein–peptide complexes. Nature Machine Intelligence 2025. DOI: 10.1038/s42256-025-01136-1.

(6) Siwek, J. C.; Omelchenko, A. A.; Chhibbar, P.; Arshad, S.; Rosengart, A.; Nazarali, I.; Patel, A.; Nazarali, K.; Rahimikollu, J.; Tilstra, J. S.;, et al. Sliding Window Interaction Grammar (SWING): a generalized interaction language model for peptide and protein interactions. Nature Methods 2025, 22 (8), 1707–1719. DOI: 10.1038/s41592-025-02723-1.

(7) Badkul, A.; Xie, L.; Zhang, S.; Xie, L. Multimodal out-of-distribution individual uncertainty quantification enhances binding affinity prediction for polypharmacology. Nature Machine Intelligence 2025, 7 (12), 1985–1995. DOI: 10.1038/s42256-025-01151-2.

(8) Huang, F.; Xia, Y.; Wang, Z.; Yang, L.; Zhang, W. Geometric Heterogeneous Graph Neural Network for Protein-Ligand Binding Affinity Prediction. In Proceedings of the 34th ACM International Conference on Information and Knowledge Management, 2025; pp 939–948.

(9) Singh, R.; Barsainyan, A. A.; Irfan, R.; Amorin, C. J.; He, S.; Davis, T.; Thiagarajan, A.; Sankaran, S.; Chithrananda, S.; Ahmad, W. ChemBERTa-3: An Open Source Training Framework for Chemical Foundation Models. ChemRxiv 2025. DOI: 10.26434/chemrxiv-2025-4glrl-v2.

(10) Ahmad, W.; Simon, E.; Chithrananda, S.; Grand, G.; Ramsundar, B. Chemberta-2: Towards chemical foundation models. arXiv preprint arXiv:2209.01712 2022.

(11) Lin, Z.; Akin, H.; Rao, R.; Hie, B.; Zhu, Z.; Lu, W.; Smetanin, N.; Verkuil, R.; Kabeli, O.; Shmueli, Y.;, et al. Evolutionary-scale prediction of atomic-level protein structure with a language model. Science 2023, 379 (6637), 1123–1130. DOI: 10.1126/science.ade2574.

(12) Hayes, T.; Rao, R.; Akin, H.; Sofroniew, N. J.; Oktay, D.; Lin, Z.; Verkuil, R.; Tran, V. Q.; Deaton, J.; Wiggert, M.;, et al. Simulating 500 million years of evolution with a language model. Science 2025, 387 (6736), 850–858. DOI: 10.1126/science.ads0018.

(13) Ji, X.; Wang, Z.; Gao, Z.; Zheng, H.; Zhang, L.; Ke, G. Uni-mol2: Exploring molecular pretraining model at scale. arXiv preprint arXiv:2406.14969 2024.

(14) Li, M.; Tan, Y.; Ma, X.; Zhong, B.; Yu, H.; Zhou, Z.; Ouyang, W.; Zhou, B.; Tan, P.; Hong, L. Prosst: Protein language modeling with quantized structure and disentangled attention. Advances in Neural Information Processing Systems 2024, 37, 35700–35726.

(15) Su, J.; He, Y.; You, S.; Jiang, S.; Zhou, X.; Zhang, X.; Wang, Y.; Su, X.; Tolstoy, I.; Chang, X.;, et al. A trimodal protein language model enables advanced protein searches. Nature Biotechnology 2025. DOI: 10.1038/s41587-025-02836-0.

(16) Ross, J.; Belgodere, B.; Chenthamarakshan, V.; Padhi, I.; Mroueh, Y.; Das, P. Large-scale chemical language representations capture molecular structure and properties. Nature Machine Intelligence 2022, 4 (12), 1256–1264. DOI: 10.1038/s42256-022-00580-7.

(17) Wang, L.; Wang, S.; Yang, H.; Li, S.; Wang, X.; Zhou, Y.; Tian, S.; Liu, L.; Bai, F. Conformational Space Profiling Enhances Generic Molecular Representation for AI-Powered Ligand-Based Drug Discovery. Advanced Science 2024, 11 (40). DOI: 10.1002/advs.202403998.

(18) Hartout, P.; Chen, D.; Pellizzoni, P.; Oliver, C.; Borgwardt, K. Endowing protein language models with structural knowledge. Bioinformatics 2025, 41 (11). DOI: 10.1093/bioinformatics/btaf582.

(19) Cheng, Z.; Xiang, H.; Ma, P.; Zeng, L.; Jin, X.; Yang, X.; Lin, J.; Feng, X.; Deng, Y.; Deng, C.;, et al. MaskMol: knowledge-guided molecular image pre-training framework for activity cliffs with pixel masking. BMC Biology 2025, 23 (1). DOI: 10.1186/s12915-025-02389-3.

(20) Zhang, Z.; Zhou, X.; Qi, Y.; Zhu, X.; Deng, X.; Tan, F.; Huang, Y.; Hu, L.; You, Z.; Hu, P. Leveraging 3D Molecular Spatial Visual Information and Multi-Perspective Representations for Drug Discovery. Advanced Science 2025. DOI: 10.1002/advs.202512453.

(21) Chen, G.; Liao, J.; Yu, Y.; Le, K.; Zhao, H.; Qin, Y.; Cai, L.; Sheng, R. HitScreen: A Sequence-Based Drug Virtual Screening Approach Using Data Augmentation and Protein Language Models. Journal of Chemical Information and Modeling 2025. DOI: 10.1021/acs.jcim.5c01753.

(22) Xu, J.; Guo, Z.; He, J.; Hu, H.; He, T.; Bai, S.; Chen, K.; Wang, J.; Fan, Y.; Dang, K. Qwen2. 5-omni technical report. arXiv preprint arXiv:2503.20215 2025.

(23) Mu, S.; Lin, S. A comprehensive survey of mixture-of-experts: Algorithms, theory, and applications. arXiv preprint arXiv:2503.07137 2025.

(24) Liu, A.; Feng, B.; Xue, B.; Wang, B.; Wu, B.; Lu, C.; Zhao, C.; Deng, C.; Zhang, C.; Ruan, C. Deepseek-v3 technical report. arXiv preprint arXiv:2412.19437 2024.

(25) Fedus, W.; Zoph, B.; Shazeer, N. Switch transformers: Scaling to trillion parameter models with simple and efficient sparsity. Journal of Machine Learning Research 2022, 23 (120), 1–39.

(26) Su, M.; Yang, Q.; Du, Y.; Feng, G.; Liu, Z.; Li, Y.; Wang, R. Comparative Assessment of Scoring Functions: The CASF-2016 Update. Journal of Chemical Information and Modeling 2019, 59 (2), 895–913. DOI: 10.1021/acs.jcim.8b00545.

(27) Stein, R. M.; Yang, Y.; Balius, T. E.; O’Meara, M. J.; Lyu, J.; Young, J.; Tang, K.; Shoichet, B. K.; Irwin, J. J. Property-unmatched decoys in docking benchmarks. Journal of chemical information and modeling 2021, 61 (2), 699–714.

(28) Tran-Nguyen, V.-K.; Jacquemard, C.; Rognan, D. LIT-PCBA: An Unbiased Data Set for Machine Learning and Virtual Screening. Journal of Chemical Information and Modeling 2020, 60 (9), 4263–4273. DOI: 10.1021/acs.jcim.0c00155.

(29) Schrodinger, LLC. The PyMOL Molecular Graphics System, Version 3.0.4. 2015.

(30) Su, J.; Han, C.; Zhou, Y.; Shan, J.; Zhou, X.; Yuan, F. SaProt: Protein Language Modeling with Structure-aware Vocabulary. bioRxiv 2024, 2023.2010.2001.560349. DOI: 10.1101/2023.10.01.560349.

(31) Fan, D.; Messmer, B.; Jaggi, M. Towards an empirical understanding of moe design choices. arXiv preprint arXiv:2402.13089 2024.

(32) Zhang, X.; Gao, H.; Wang, H.; Chen, Z.; Zhang, Z.; Chen, X.; Li, Y.; Qi, Y.; Wang, R. PLANET: A Multi-objective Graph Neural Network Model for Protein–Ligand Binding Affinity Prediction. Journal of Chemical Information and Modeling 2023, 64 (7), 2205–2220. DOI: 10.1021/acs.jcim.3c00253.

(33) Huang, A.; Knight, I. S.; Naprienko, S. Data leakage and redundancy in the lit-pcba benchmark. arXiv preprint arXiv:2507.21404 2025.

(34) Wang, R.; Fang, X.; Lu, Y.; Wang, S. The PDBbind Database: Collection of Binding Affinities for Protein-Ligand Complexes with Known Three-Dimensional Structures. Journal of Medicinal Chemistry 2004, 47 (12), 2977–2980. DOI: 10.1021/jm030580l.

(35) Liu, Z.; Li, Y.; Han, L.; Li, J.; Liu, J.; Zhao, Z.; Nie, W.; Liu, Y.; Wang, R. PDB-wide collection of binding data: current status of the PDBbind database. Bioinformatics 2014, 31 (3), 405–412. DOI: 10.1093/bioinformatics/btu626.

(36) Zhang, Y.; Skolnick, J. Scoring function for automated assessment of protein structure template quality. Proteins: Structure, Function, and Bioinformatics 2004, 57 (4), 702–710. DOI: 10.1002/prot.20264 (acccessed 2026/03/13).

37. (37) Wang, L.; Pulugurta, R.; Vure, P.; Zhang, Y.; Pal, A.; Chatterjee, P. Pepdora: A unified peptide language model via weight-decomposed low-rank adaptation. arXiv preprint arXiv:2410.20667 2024.

(38) Yang, A.; Li, A.; Yang, B.; Zhang, B.; Hui, B.; Zheng, B.; Yu, B.; Gao, C.; Huang, C.; Lv, C. Qwen3 technical report. arXiv preprint arXiv:2505.09388 2025.

(39) Team, K.; Bai, Y.; Bao, Y.; Charles, Y.; Chen, C.; Chen, G.; Chen, H.; Chen, H.; Chen, J.; Chen, N. Kimi k2: Open agentic intelligence. arXiv preprint arXiv:2507.20534 2025.

(40) Graber, D.; Stockinger, P.; Meyer, F.; Mishra, S.; Horn, C.; Buller, R. Resolving data bias improves generalization in binding affinity prediction. Nature Machine Intelligence 2025. DOI: 10.1038/s42256-025-01124-5.

(41) Xu, S.; Shen, L.; Zhang, M.; Jiang, C.; Zhang, X.; Xu, Y.; Liu, J.; Liu, X. Surface-based multimodal protein–ligand binding affinity prediction. Bioinformatics 2024, 40 (7). DOI: 10.1093/bioinformatics/btae413.

(42) Mukta, F. T.; Rana, M. M.; Meyer, A.; Ellingson, S.; Nguyen, D. D. The algebraic extended atom-type graph-based model for precise ligand–receptor binding affinity prediction. Journal of Cheminformatics 2025, 17 (1). DOI: 10.1186/s13321-025-00955-z.

(43) Dong, L.; Shi, S.; Qu, X.; Luo, D.; Wang, B.; Dong, L. Ligand binding affinity prediction with fusion of graph neural networks and 3D structure-based complex graph. Physical Chemistry Chemical Physics 2023, 25 (35), 24110–24120, 10.1039/D3CP03651K. DOI: 10.1039/D3CP03651K.

(44) Min, Y.; Wei, Y.; Wang, P.; Wang, X.; Li, H.; Wu, N.; Bauer, S.; Zheng, S.; Shi, Y.; Wang, Y.;, et al. From Static to Dynamic Structures: Improving Binding Affinity Prediction with Graph-Based Deep Learning. Advanced Science 2024, 11 (40). DOI: 10.1002/advs.202405404.

(45) Hu, F.; Jiang, J.; Wang, D.; Zhu, M.; Yin, P. Multi-PLI: interpretable multi-task deep learning model for unifying protein–ligand interaction datasets. Journal of Cheminformatics 2021, 13 (1). DOI: 10.1186/s13321-021-00510-6.

(46) Wang, K.; Song, A.; Liu, F.; Luan, X.; Wang, X.; Zhou, J. PLMAM-PLA: A Method Using Pretrained Language Models and Attention Mechanisms for Protein-Ligand Binding Affinity Prediction. IEEE Transactions on Computational Biology and Bioinformatics 2025, 1–12. DOI: 10.1109/TCBBIO.2025.3583738.

(47) Davis, M. I.; Hunt, J. P.; Herrgard, S.; Ciceri, P.; Wodicka, L. M.; Pallares, G.; Hocker, M.; Treiber, D. K.; Zarrinkar, P. P. Comprehensive analysis of kinase inhibitor selectivity. Nature Biotechnology 2011, 29 (11), 1046–1051. DOI: 10.1038/nbt.1990.

(48) Tang, J.; Szwajda, A.; Shakyawar, S.; Xu, T.; Hintsanen, P.; Wennerberg, K.; Aittokallio, T. Making Sense of Large-Scale Kinase Inhibitor Bioactivity Data Sets: A Comparative and Integrative Analysis. Journal of Chemical Information and Modeling 2014, 54 (3), 735–743. DOI: 10.1021/ci400709d.

(49) Liu, T.; Lin, Y.; Wen, X.; Jorissen, R. N.; Gilson, M. K. BindingDB: a web-accessible database of experimentally determined protein-ligand binding affinities. Nucleic Acids Research 2007, 35 (Database), D198–D201. DOI: 10.1093/nar/gkl999.

(50) Shah, P. M.; Zhu, H.; Lu, Z.; Wang, K.; Tang, J.; Li, M. DeepDTAGen: a multitask deep learning framework for drug-target affinity prediction and target-aware drugs generation. Nature Communications 2025, 16 (1). DOI: 10.1038/s41467-025-59917-6.

(51) Chen, Y.; Huang, J.; Liu, C.; Zhang, S.; Li, X.; Zhang, Z.; Chen, T. G.; Wang, L. DualPG-DTA: A Large Language Model-Powered Graph Neural Network Framework for Enhanced Drug-Target Affinity Prediction and Discovery of Novel CDK9 Inhibitors Exhibiting In Vivo Anti-Leukemia Activity. Advanced Science 2026, 13 (12). DOI: 10.1002/advs.202513099.

(52) Bidgoli, A. H.; Mahdavi, M.; Malek, H. Structure-free drug–target affinity prediction using protein and molecule language models. Journal of Cheminformatics 2026, 18 (1), 21. DOI: 10.1186/s13321-025-01146-6.

(53) Eddy, S. R. Accelerated Profile HMM Searches. PLOS Computational Biology 2011, 7 (10), e1002195. DOI: 10.1371/journal.pcbi.1002195.

(54) Mistry, J.; Chuguransky, S.; Williams, L.; Qureshi, M.; Salazar, Gustavo A.; Sonnhammer, E. L. L.; Tosatto, S. C. E.; Paladin, L.; Raj, S.; Richardson, L. J.;, et al. Pfam: The protein families database in 2021. Nucleic Acids Research 2021, 49 (D1), D412–D419. DOI: 10.1093/nar/gkaa913 (acccessed 3/14/2026).

(55) Mahdizadeh, S. J.; Eriksson, L. A. MolAI: A Deep Learning Framework for Data-Driven Molecular Descriptor Generation and Advanced Drug Discovery Applications. Journal of Chemical Information and Modeling 2025. DOI: 10.1021/acs.jcim.5c00491.

(56) Yüksel, A.; Ulusoy, E.; Ünlü, A.; Doğan, T. SELFormer: molecular representation learning via SELFIES language models. Machine Learning: Science and Technology 2023, 4 (2), 025035. DOI: 10.1088/2632-2153/acdb30.

(57) Zeng, X.; Xiang, H.; Yu, L.; Wang, J.; Li, K.; Nussinov, R.; Cheng, F. Accurate prediction of molecular properties and drug targets using a self-supervised image representation learning framework. Nature Machine Intelligence 2022, 4 (11), 1004–1016. DOI: 10.1038/s42256-022-00557-6.

## Reference

(1) Ahmad, W.; Simon, E.; Chithrananda, S.; Grand, G.; Ramsundar, B. Chemberta-2: Towards chemical foundation models. arXiv preprint arXiv:2209.01712 2022.

(2) Singh, R.; Barsainyan, A. A.; Irfan, R.; Amorin, C. J.; He, S.; Davis, T.; Thiagarajan, A.; Sankaran, S.; Chithrananda, S.; Ahmad, W. ChemBERTa-3: An Open Source Training Framework for Chemical Foundation Models. ChemRxiv 2025. DOI: 10.26434/chemrxiv-2025-4glrl-v2.

(3) Wang, L.; Pulugurta, R.; Vure, P.; Zhang, Y.; Pal, A.; Chatterjee, P. PepDoRA: A Unified Peptide Language Model via Weight-Decomposed Low-Rank Adaptation. 2024.

(4) Ross, J.; Belgodere, B.; Chenthamarakshan, V.; Padhi, I.; Mroueh, Y.; Das, P. Large-scale chemical language representations capture molecular structure and properties. Nature Machine Intelligence 2022, 4 (12), 1256–1264. DOI: 10.1038/s42256-022-00580-7.

(5) Mahdizadeh, S. J.; Eriksson, L. A. MolAI: A Deep Learning Framework for Data-Driven Molecular Descriptor Generation and Advanced Drug Discovery Applications. Journal of Chemical Information and Modeling 2025. DOI: 10.1021/acs.jcim.5c00491.

(6) Yüksel, A.; Ulusoy, E.; Ünlü, A.; Doğan, T. SELFormer: molecular representation learning via SELFIES language models. Machine Learning: Science and Technology 2023, 4 (2), 025035. DOI: 10.1088/2632-2153/acdb30.

(7) Wang, L.; Wang, S.; Yang, H.; Li, S.; Wang, X.; Zhou, Y.; Tian, S.; Liu, L.; Bai, F. Conformational Space Profiling Enhances Generic Molecular Representation for AI-Powered Ligand-Based Drug Discovery. Advanced Science 2024, 11 (40). DOI: 10.1002/advs.202403998.

(8) Ji, X.; Wang, Z.; Gao, Z.; Zheng, H.; Zhang, L.; Ke, G. Uni-mol2: Exploring molecular pretraining model at scale. arXiv preprint arXiv:2406.14969 2024.

(9) Zeng, X.; Xiang, H.; Yu, L.; Wang, J.; Li, K.; Nussinov, R.; Cheng, F. Accurate prediction of molecular properties and drug targets using a self-supervised image representation learning framework. Nature Machine Intelligence 2022, 4 (11), 1004–1016. DOI: 10.1038/s42256-022-00557-6.

(10) Cheng, Z.; Xiang, H.; Ma, P.; Zeng, L.; Jin, X.; Yang, X.; Lin, J.; Feng, X.; Deng, Y.; Deng, C.;, et al. MaskMol: knowledge-guided molecular image pre-training framework for activity cliffs with pixel masking. BMC Biology 2025, 23 (1). DOI: 10.1186/s12915-025-02389-3.

(13) Su, J.; Han, C.; Zhou, Y.; Shan, J.; Zhou, X.; Yuan, F. SaProt: Protein Language Modeling with Structure-aware Vocabulary. bioRxiv 2024, 2023.2010.2001.560349. DOI: 10.1101/2023.10.01.560349.

(15) Yang, Z.; Zhong, W.; Lv, Q.; Dong, T.; Chen, G.; Chen, C. Y.-C. Interaction-Based Inductive Bias in Graph Neural Networks: Enhancing Protein-Ligand Binding Affinity Predictions From 3D Structures. IEEE Transactions on Pattern Analysis and Machine Intelligence 2024, 1–18. DOI: 10.1109/TPAMI.2024.3400515.

(16) Wang, L.; Pulugurta, R.; Vure, P.; Zhang, Y.; Pal, A.; Chatterjee, P. Pepdora: A unified peptide language model via weight-decomposed low-rank adaptation. arXiv preprint arXiv:2410.20667 2024.

(17) He, K.; Zhang, X.; Ren, S.; Sun, J. Deep residual learning for image recognition. In Proceedings of the IEEE conference on computer vision and pattern recognition, 2016; pp 770–778.

(18) Dosovitskiy, A.; Beyer, L.; Kolesnikov, A.; Weissenborn, D.; Zhai, X.; Unterthiner, T.; Dehghani, M.; Minderer, M.; Heigold, G.; Gelly, S. An image is worth 16×16 words: Transformers for image recognition at scale. arXiv preprint arXiv:2010.11929 2020.

(19) Schrodinger, LLC. The PyMOL Molecular Graphics System, Version 3.0.0. In The PyMOL Molecular Graphics System, Version 3.0.0, 2025.

(20) Eddy, S. R. Accelerated Profile HMM Searches. PLOS Computational Biology 2011, 7 (10), e1002195. DOI: 10.1371/journal.pcbi.1002195.

(21) Mistry, J.; Chuguransky, S.; Williams, L.; Qureshi, M.; Salazar, Gustavo A.; Sonnhammer, E. L. L.; Tosatto, S. C. E.; Paladin, L.; Raj, S.; Richardson, L. J.;, et al. Pfam: The protein families database in 2021. Nucleic Acids Research 2021, 49 (D1), D412–D419. DOI: 10.1093/nar/gkaa913 (acccessed 3/14/2026).

(22) Su, M.; Yang, Q.; Du, Y.; Feng, G.; Liu, Z.; Li, Y.; Wang, R. Comparative Assessment of Scoring Functions: The CASF-2016 Update. Journal of Chemical Information and Modeling 2019, 59 (2), 895–913. DOI: 10.1021/acs.jcim.8b00545.

(23) Zhang, Y.; Skolnick, J. Scoring function for automated assessment of protein structure template quality. Proteins: Structure, Function, and Bioinformatics 2004, 57 (4), 702–710. DOI: 10.1002/prot.20264 (acccessed 2026/03/13).

(24) Graber, D.; Stockinger, P.; Meyer, F.; Mishra, S.; Horn, C.; Buller, R. Resolving data bias improves generalization in binding affinity prediction. Nature Machine Intelligence 2025. DOI: 10.1038/s42256-025-01124-5.

(25) Shen, C.; Zhang, X.; Hsieh, C.-Y.; Deng, Y.; Wang, D.; Xu, L.; Wu, J.; Li, D.; Kang, Y.; Hou, T.;, et al. A generalized protein–ligand scoring framework with balanced scoring, docking, ranking and screening powers. Chemical Science 2023, 14 (30), 8129–8146, 10.1039/D3SC02044D. DOI: 10.1039/D3SC02044D.

(26) Stepniewska-Dziubinska, M. M.; Zielenkiewicz, P.; Siedlecki, P. Development and evaluation of a deep learning model for protein–ligand binding affinity prediction. Bioinformatics 2018, 34 (21), 3666–3674. DOI: 10.1093/bioinformatics/bty374.

(27) Xu, S.; Shen, L.; Zhang, M.; Jiang, C.; Zhang, X.; Xu, Y.; Liu, J.; Liu, X. Surface-based multimodal protein–ligand binding affinity prediction. Bioinformatics 2024, 40 (7). DOI: 10.1093/bioinformatics/btae413.

(28) Mukta, F. T.; Rana, M. M.; Meyer, A.; Ellingson, S.; Nguyen, D. D. The algebraic extended atom-type graph-based model for precise ligand–receptor binding affinity prediction. Journal of Cheminformatics 2025, 17 (1). DOI: 10.1186/s13321-025-00955-z.

(29) Dong, L.; Shi, S.; Qu, X.; Luo, D.; Wang, B.; Dong, L. Ligand binding affinity prediction with fusion of graph neural networks and 3D structure-based complex graph. Physical Chemistry Chemical Physics 2023, 25 (35), 24110–24120, 10.1039/D3CP03651K. DOI: 10.1039/D3CP03651K.

(30) Min, Y.; Wei, Y.; Wang, P.; Wang, X.; Li, H.; Wu, N.; Bauer, S.; Zheng, S.; Shi, Y.; Wang, Y.;, et al. From Static to Dynamic Structures: Improving Binding Affinity Prediction with Graph-Based Deep Learning. Advanced Science 2024, 11 (40). DOI: 10.1002/advs.202405404.

(31) Hu, F.; Jiang, J.; Wang, D.; Zhu, M.; Yin, P. Multi-PLI: interpretable multi-task deep learning model for unifying protein–ligand interaction datasets. Journal of Cheminformatics 2021, 13 (1). DOI: 10.1186/s13321-021-00510-6.

(32) Wang, K.; Song, A.; Liu, F.; Luan, X.; Wang, X.; Zhou, J. PLMAM-PLA: A Method Using Pretrained Language Models and Attention Mechanisms for Protein-Ligand Binding Affinity Prediction. IEEE Transactions on Computational Biology and Bioinformatics 2025, 1–12. DOI: 10.1109/TCBBIO.2025.3583738.

(33) Davis, M. I.; Hunt, J. P.; Herrgard, S.; Ciceri, P.; Wodicka, L. M.; Pallares, G.; Hocker, M.; Treiber, D. K.; Zarrinkar, P. P. Comprehensive analysis of kinase inhibitor selectivity. Nature Biotechnology 2011, 29 (11), 1046–1051. DOI: 10.1038/nbt.1990.

(34) Tang, J.; Szwajda, A.; Shakyawar, S.; Xu, T.; Hintsanen, P.; Wennerberg, K.; Aittokallio, T. Making Sense of Large-Scale Kinase Inhibitor Bioactivity Data Sets: A Comparative and Integrative Analysis. Journal of Chemical Information and Modeling 2014, 54 (3), 735–743. DOI: 10.1021/ci400709d.

(35) Liu, T.; Lin, Y.; Wen, X.; Jorissen, R. N.; Gilson, M. K. BindingDB: a web-accessible database of experimentally determined protein-ligand binding affinities. Nucleic Acids Research 2007, 35 (Database), D198–D201. DOI: 10.1093/nar/gkl999.

(36) Shah, P. M.; Zhu, H.; Lu, Z.; Wang, K.; Tang, J.; Li, M. DeepDTAGen: a multitask deep learning framework for drug-target affinity prediction and target-aware drugs generation. Nature Communications 2025, 16 (1). DOI: 10.1038/s41467-025-59917-6.

(37) Chen, Y.; Huang, J.; Liu, C.; Zhang, S.; Li, X.; Zhang, Z.; Chen, T. G.; Wang, L. DualPG-DTA: A Large Language Model-Powered Graph Neural Network Framework for Enhanced Drug-Target Affinity Prediction and Discovery of Novel CDK9 Inhibitors Exhibiting In Vivo Anti-Leukemia Activity. Advanced Science 2026, 13 (12). DOI: 10.1002/advs.202513099.

(38) Bidgoli, A. H.; Mahdavi, M.; Malek, H. Structure-free drug–target affinity prediction using protein and molecule language models. Journal of Cheminformatics 2026, 18 (1), 21. DOI: 10.1186/s13321-025-01146-6.

(39) Öztürk, H.; Özgür, A.; Ozkirimli, E. DeepDTA: deep drug–target binding affinity prediction. Bioinformatics 2018, 34 (17), i821–i829. DOI: 10.1093/bioinformatics/bty593.

(40) Zhang, L.; Zeng, W.; Chen, J.; Chen, J.; Li, K. GDilatedDTA: Graph dilation convolution strategy for drug target binding affinity prediction. Biomedical Signal Processing and Control 2024, 92, 106110. DOI: 10.1016/j.bspc.2024.106110.

(41) Shah, P. M.; Zhu, H.; Lu, Z.; Wang, K.; Tang, J.; Li, M. DeepDTAGen: a multitask deep learning framework for drug-target affinity prediction and target-aware drugs generation. Nature Communications 2025, 16 (1), 5021. DOI: 10.1038/s41467-025-59917-6.

